# Inter-domain dynamics in the chaperone SurA and multi-site binding to its unfolded outer membrane protein clients

**DOI:** 10.1101/2019.12.19.882696

**Authors:** Antonio N. Calabrese, Bob Schiffrin, Matthew Watson, Theodoros K. Karamanos, Martin Walko, Julia R. Humes, Jim E. Horne, Paul White, Andrew J. Wilson, Antreas C. Kalli, Roman Tuma, Alison E. Ashcroft, David J. Brockwell, Sheena E. Radford

## Abstract

The periplasmic chaperone SurA plays a key role in outer membrane protein (OMP) biogenesis. *E. coli* SurA comprises a core domain and two peptidylprolyl isomerase domains (P1 and P2), but how it binds its OMP clients and the mechanism(s) of its chaperone action remain unclear. Here, we have used chemical cross-linking, hydrogen-deuterium exchange, single-molecule FRET and molecular dynamics simulations to map the client binding site(s) on SurA and to interrogate the role of conformational dynamics of the chaperone’s domains in OMP recognition. We demonstrate that SurA samples a broad array of conformations in solution in which P2 primarily lies closer to the core/P1 domains than suggested by its crystal structure. Multiple binding sites for OMPs are located primarily in the core domain, with binding of the unfolded OMP resulting in conformational changes between the core/P1 domains. Together, the results portray a model in which unfolded OMP substrates bind in a cradle formed between the SurA domains, with structural flexibility between its domains assisting OMP recognition, binding and release.

## Introduction

Chaperones play vital roles in multicomponent proteostasis networks, ensuring that proteins fold and avoid aggregation in the crowded cellular milieu, and that misfolded proteins which cannot be rescued by chaperones are targeted for degradation ^1, 2^. It is now established that many chaperones are in rapid dynamic exchange between co-populated conformations, and that this conformational plasticity is key to their functional mechanisms ^3^. In the case of ATP-dependent chaperones, e.g. the Hsp60 chaperonins GroEL and TRiC, and the Hsp90 and Hsp70 families, ATP binding and/or hydrolysis promotes conformational changes that facilitate the folding and/or release of their clients ^1, 2, 4, 5, 6, 7, 8^. However, some chaperones are not dependent on energy from nucleotide binding/hydrolysis, and instead their intrinsic structural flexibility is proposed to be key to their function ^3, 9, 10, 11, 12^. The functional mechanisms of these ATP-independent chaperones, including how they bind and release their substrates in a controlled manner, is generally not well understood.

SurA is an ATP-independent chaperone involved in the biogenesis of outer membrane proteins (OMPs) in the periplasm of Gram-negative bacteria ^13 – 18^. This protein is thought to be the major chaperone responsible for protecting OMPs from aggregation in the periplasm ^13 – 18^ and facilitating OMP delivery to the β-barrel assembly machinery (BAM) for folding and insertion into the outer membrane (OM) ^13, 14, 19, 20, 21^. Deletion of SurA leads to OMP assembly defects, the induction of stress responses, and increased sensitivity to antibiotics and detergents ^15, 17, 22, 23, 24^. Further, *ΔsurA* strains show reduced assembly of virulence factors, such as pili and adhesins, and exhibit reduced pathogenicity in a number of species ^22, 25, 26^. *E. coli* SurA has a three domain architecture, consisting of a core domain which is composed of its N- and C-terminal regions, and two parvulin-like peptidylprolyl isomerase (PPIase) domains (P1 and P2) (Fig. 1a) ^27^. However, despite the availability of its crystal structure [27], how SurA binds its unfolded OMP clients and the molecular mechanism(s) of SurA function remain unknown. A substrate binding crevice was proposed based on examination of molecular packing interactions in crystals of SurA (Fig. 1b), but the location of OMP binding regions and the roles of the PPIase domains (which are not essential for *in vivo* or *in vitro* function, at least for some clients ^24, 28, 29^) in folding and binding its varied OMP clients remained unknown.

**Fig. 1.**
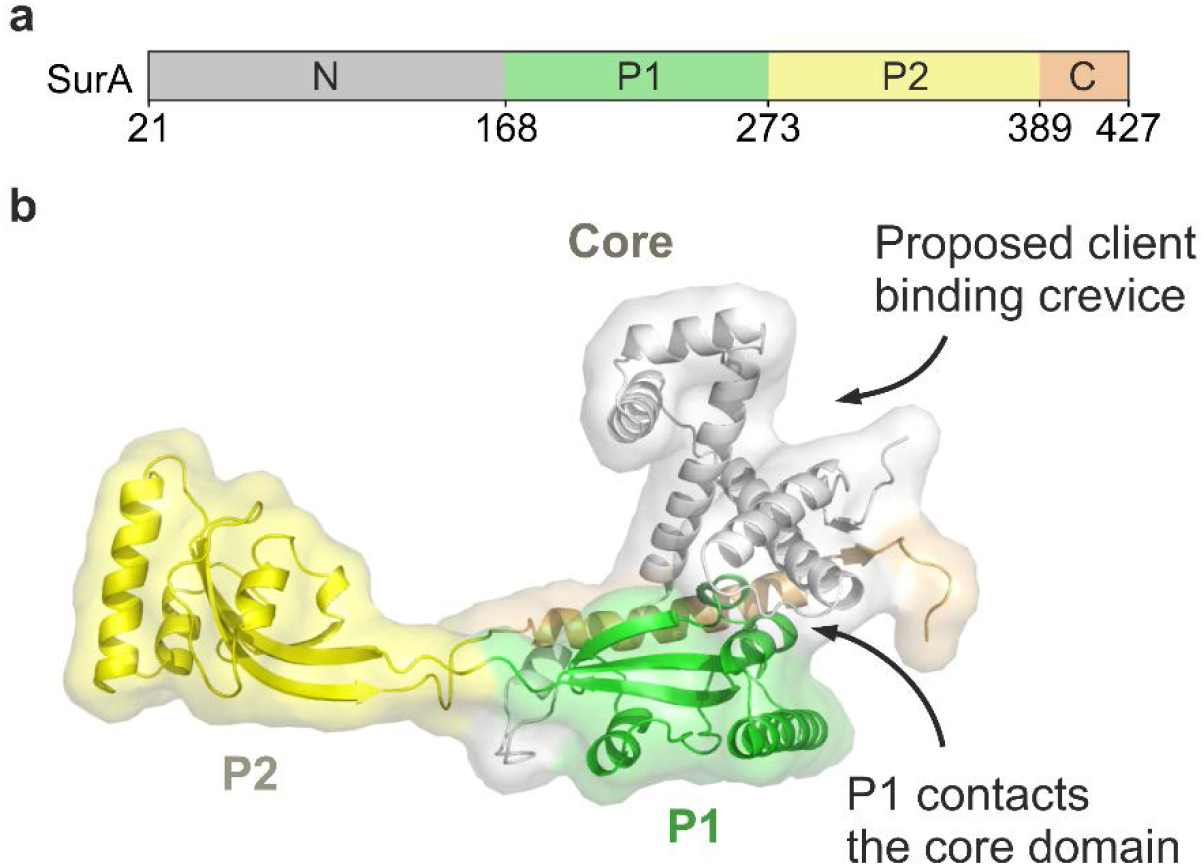
SurA structure and domain architecture. **(a)** Domain architecture of *E. coli* SurA. Regions are coloured grey (N-terminal region of the core domain), green (P1), yellow (P2) and orange (C-terminal region of the core domain). The signal sequence is not shown, and was not present in any of the constructs used in this study, but the numbering used throughout reflects the gene numbering (including the signal peptide). **(b)** Crystal structure of *E. coli* SurA WT (PDB: 1M5Y ^27^), with missing residues added using MODELLER [51]. A client binding crevice in the core domain was proposed based on crystal contacts with a neighbouring SurA molecule, as indicated ^27^. P1 contacts the core domain in this crystal structure (**Supplementary Fig. 1a-c**). Regions are coloured as in **(a)**.

In the crystal structure of full-length SurA ^27^, an extended conformation is observed in which the core and P1 domains are in contact (Fig. 1b, **Supplementary Fig. 1a-c**), while P2 is separated from this globular region via a linker (Fig. 1b, **Supplementary Fig. 1d**). Examination of the molecular packing interactions in the crystal lattice revealed multiple contacts between all three domains and neighbouring molecules, which may stabilise the elongated architecture observed (**Supplementary Fig. 2**). SurA homologues and domain deletion variants have been crystallised in conformations with a variety of domain orientations (**Supplementary Fig. 3**) suggesting that SurA may have a dynamic structure. Further, tethering of the P1 and core domains via a disulfide bond resulted in impaired OMP assembly *in vivo* ^29^. However, the precise nature of these conformational dynamics and how they are linked to OMP binding have remained elusive.

Here, we sought to determine the conformational properties of full-length *E. coli* SurA in solution in an effort to better understand its conformational dynamics and how inter-domain motions may be exploited or modified by client binding. Combining mass spectrometric (MS) methods (chemical crosslinking (XL) and hydrogen-deuterium exchange (HDX)), with single-molecule FRET (smFRET) and molecular dynamics (MD) simulations we show that SurA adopts conformations in solution that differ substantially from its crystal structure ^27^. Specifically, the P1 domain samples open and closed states relative to the core domain, and P2 is primarily located closer to the core/P1 domains than observed in the crystal structure. We also show that multiple sites on the SurA surface, predominantly located in the core domain, are involved in client binding. OMPs bind to these specific sites in different orientations, consistent with a dynamically bound state, and the conformations adopted by the chaperone alter in response to OMP binding. Combined, our results portray a model in which the three domains of SurA form a cradle around its OMP clients, protecting them from misfolding and aggregation on their journey through the periplasm, with the conformational dynamics of the domains presumably facilitating their delivery to BAM for folding into the outer membrane.

## Results

### Inter-domain conformational flexibility in SurA

We first investigated the structure and dynamics of apo-SurA in solution using XL-MS, which provides distance information in the form of spatial restraints, and enables comparison of the solution conformation(s) of the protein with structural data ^30^. For this purpose, we used the bifunctional reagent disuccinimidyl dibutyric urea (DSBU), which primarily crosslinks Lys residues ^31^. DSBU has been shown to crosslink residues within a straight line distance (SLD) between their Cα atoms of *ca*. 27-30 Å ^32^. More recently it has been shown that considering the solvent accessible surface distance (SASD) between residues may more reliably predict structural models ^33^ (a Cα− Cα distance between crosslinked residues of up to *ca.* 35 Å is considered feasible for DSBU). For monomeric SurA, a total of 13 intra-domain (core-core, P1-P1 and P2-P2) (**Supplementary Fig. 4, Supplementary Table 1**) and 19 inter-domain (core-P1, core-P2 and P1-P2) crosslinks were detected (Fig. 2a, **Supplementary Table 1**). Most of the intra-domain crosslinks identified (8 of 13, based on the SASD) are consistent with the domain structures observed in the crystal structure of full-length *E. coli* SurA (**Supplementary Fig. 4, Supplementary Table 1**), with the 5 unsatisfied crosslinks (K90-K134, K134-K394 and K134-K405 in the core domain, along with K278-K293 and K362-K388 in the P2 domain) being located in flexible, likely dynamic, loop regions either within, or between domains (**Supplementary Fig. 1d**). By contrast, only four (K105-K278, K278-K394, K251-K278, K252-278) of the 19 inter-domain crosslinks (core-P1, core-P2 or P1-P2) are compatible with the SurA crystal structure based on the SASD (Fig. 2b-d, **Supplementary Table 1**). Interestingly, the three core-P1 crosslinks detected (K251-K405, K252-K394 and K269-K394) involve residues that have SLDs of ~26 Å ^34^. However, we noted that the vectors of the SLDs for these crosslinks pass directly through the P1 and core domains, highlighting the importance of considering SASDs in judging crosslink violation ^33, 35^. Taken together, the data show that SurA populates structures in solution in which P2 is closer to both the core and P1 domains than portrayed by its crystal structure, as well as conformations in which the orientation and/or distance of P1 relative to the core is distinct from that observed in the crystal structure of the protein ^27^.

**Fig. 2.**
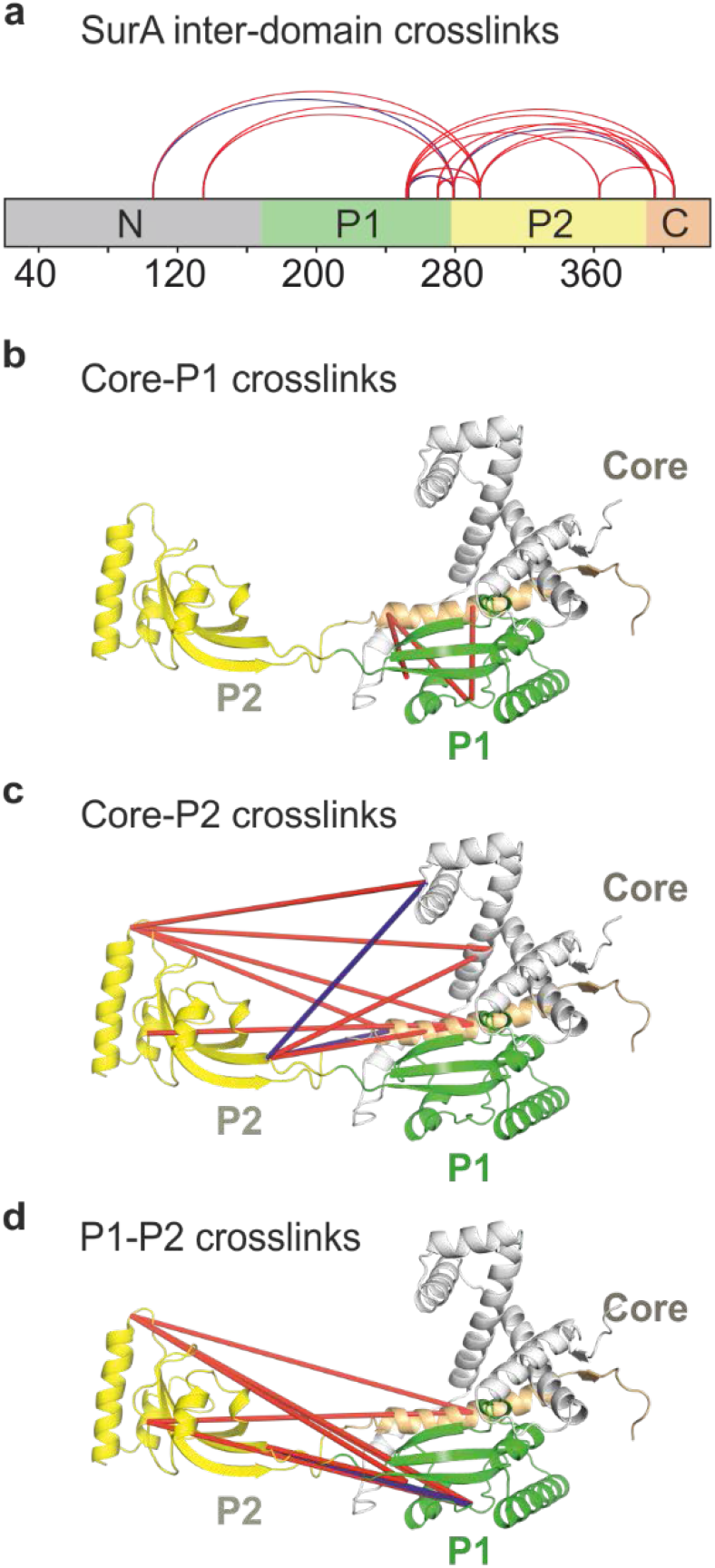
XL-MS suggests the P2 domain is closer to the core/P1 than implied by the crystal structure. **(a)** Locations of the 19 identified SurA inter-domain crosslinks (red and blue lines). **(b-d)** Crystal structure of SurA showing the identified inter-domain crosslinks between **(b)** core-P1 (3 in total), **(c)** core-P2 (9 in total), and **(d)** P1-P2 (7 in total). Only four of the inter-domain crosslinks identified (coloured in blue) are consistent with the crystal structure of full-length SurA (PDB: 1M5Y ^27^), based on a maximum SASD of 35 Å ^33^. Other crosslinks are inconsistent with this distance cut-off (red lines). Details of crosslinked residues are given in **Supplementary Table 1**. Reactions contained 5 µM SurA, 50 µM DSBU, in 10 mM potassium phosphate buffer, pH 8.0, for 45 min, 25 °C. Note that for clarity crosslinks are shown as straight lines between residues (rather than as SASD which provide a more reliable measurement for comparison with protein structures (see text)). The N-terminal region of the core domain, P1, P2, and the C-terminal region of the core domain are shown in grey, green, yellow, and orange, respectively.

As an independent validation of the crosslinking results, we investigated the inter-domain distances in SurA by smFRET ^36^. We selected non-conserved residues in the core, P1 and P2 domains (Q85, N193, and E301, respectively) to substitute with Cys, and constructed three variants containing Cys substitutions at two positions (core-P1, core-P2 and P1-P2) (Fig. 3a-c, **Supplementary Table 2**). Each SurA variant was stochastically labelled with Alexa 488 and Alexa 594 dyes (R_0_ = 60 Å), enabling inter-domain distance distributions to be monitored in a pairwise fashion. Samples containing ~50 pM of labelled SurA were interrogated using confocal fluorescence detection and alternating laser excitation (ALEX) (see Methods). Fluorescence intensities in the donor and acceptor channels yielded FRET efficiencies (E_FRET_) for the passage of each single molecule through the confocal volume (fluorescence burst). These were collated into FRET efficiency histograms and compared with distributions predicted for each labelled SurA double Cys variant calculated from the SurA crystal structure (see Methods) (Fig. 3d-f) ^37, 38^.

**Fig. 3.**
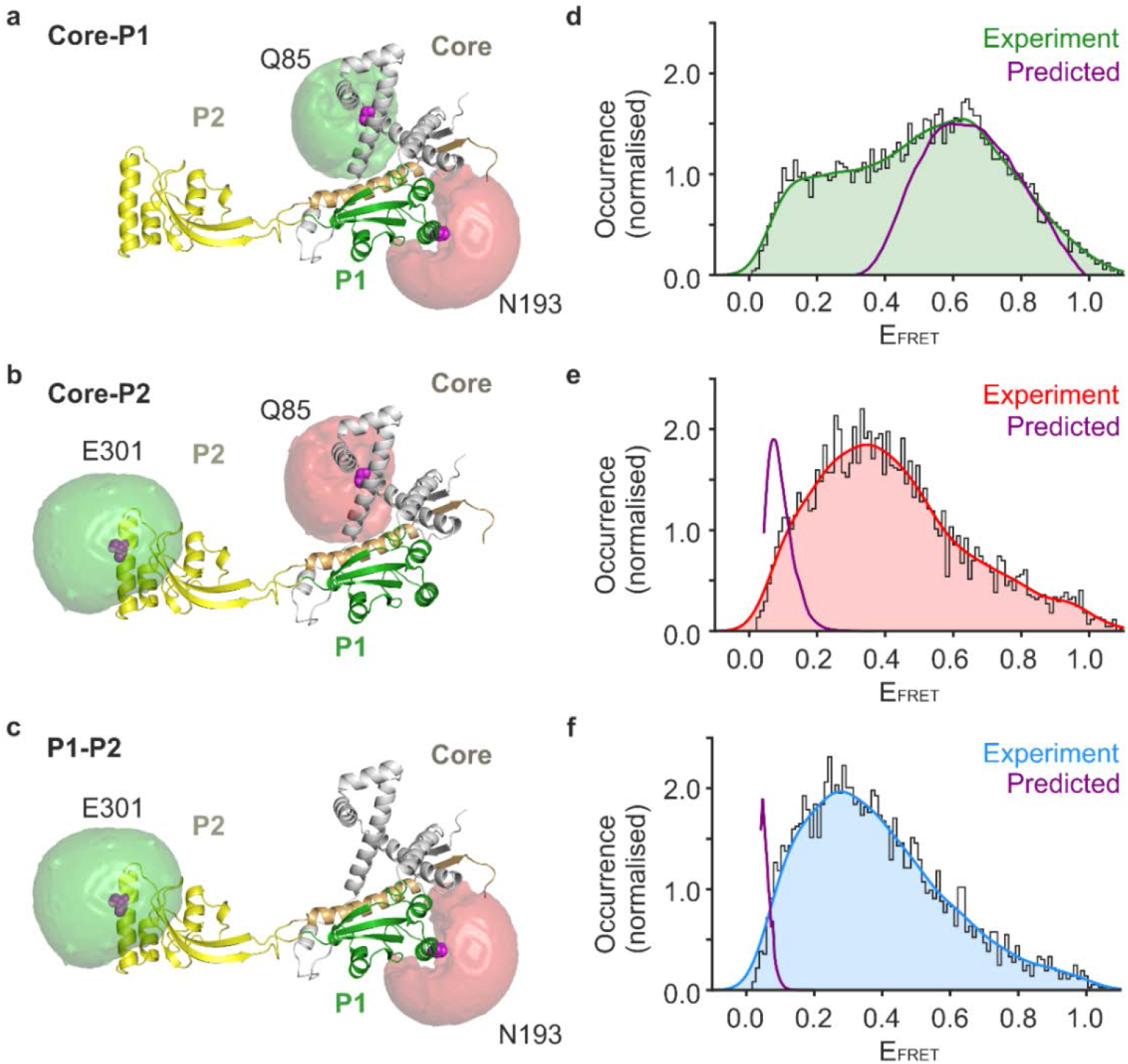
smFRET experiments confirm dramatic differences in inter-domain distances between SurA in solution and those predicted from its crystal structure. **(a-c)** Locations of residues in each SurA domain mutated to Cys for pairwise labelling with FRET dyes are shown as magenta spheres. Predicted available volumes for AlexaFluor 488 and AlexaFluor 594 dyes attached to the Cys residues (magenta) represented by green and red surfaces, respectively. Note that Cys residues were labelled stochastically and only one double-labelled example is shown. **(d-f)** Experimentally measured and predicted E_FRET_ distributions for the three pairwise combinations of fluorescently-labelled SurA double mutants. Histograms of the measured E_FRET_ values are shown in grey, and kernel density estimations (KDEs) of the probability density function of the measured E_FRET_ values are shown in green, red and blue for the core-P1, core-P2 and P1-P2 pairwise measurements, respectively (see Methods). Predicted E_FRET_ distributions (purple lines) were calculated from the SurA crystal structure (PDB: 1M5Y ^27^) using distance distributions generated by the MtsslWizard plugin for PyMOL ^87^, which takes into account both the location of the dyes and the flexibility of the dye linkers ^37^. Note that all dyes can be considered as freely rotating, as manifested by low anisotropy (**Supplementary Table 2**), hence changes in E_FRET_ can be translated to distance variations. Samples contained ~50 pM labelled SurA variant in 50 mM Tris-HCl, pH 8.0, 25 °C.

The predicted E_FRET_ distribution for the core-P1 SurA variant calculated from the crystal structure of *E. coli* SurA ^27^ has a single maximum at ~0.6 (Fig. 3d). In marked contrast with this prediction, the measured E_FRET_ distribution of the core-P1 labelled SurA is (at least) bimodal, with one population centred on E_FRET_ ~0.6, and a second, smaller, population centred on E_FRET_ ~0.2 (Fig. 3d). This suggests that SurA populates at least two distinct conformations in solution, one (~60 %) in which P1 is located close to the core domain with an inter-domain distance similar to that in the crystal structure (core-P1_closed_), and one (~40 %) in which the P1 and core domains are further apart (core-P1_open_). The observed E_FRET_ distributions for the labelled core-P2 and P1-P2 variants have maxima at ~0.4 and ~0.3, respectively (Fig. 3e,f), both in marked contrast with the low predicted values in the crystal structure (~0.1 and ~0.02, a spatial separation of ~85 Å and ~115 Å, for core-P2 and P1-P2, respectively). This indicates that, in the vast majority of molecules, P2 is located closer to the core and P1 domains than suggested by the SurA crystal structure ^27^, consistent with the XL-MS data (Fig. 2b,c,d). Burst variance analysis (BVA) ^39^ showed that inter-domain motions involving each pair of domains occurs on the timescale of diffusion through the confocal volume (<1 ms) (**Supplementary Fig. 5a-c**). Together, these data indicate a dynamic chaperone structure in which sub-ms motions involving all three domains are occurring, in particular at the core-P1 interface which interconverts between core-P1_closed_ and core-P1_open_ states. In addition, the data show that P2 spends most of its time closer to the core and P1 domains than suggested by the SurA crystal structure.

To help visualise the possible conformational excursions of the different domains of apo-SurA in solution we performed unrestrained all-atom molecular dynamics (MD) simulations. Initially three 1 μs simulations were performed starting from the crystal structure of full-length SurA (PDB: 1M5Y ^27^). However, in these simulations the individual domains of SurA were unstable, with both P1 and P2 unfolding (not shown). Therefore, we built an alternative starting model of full-length SurA in which the core and P1 domains are spatially separated, consistent with the smFRET data (SurA^core-P1-open^) (**Supplementary Fig. 6a**, see Methods). In the three 1 μs simulations performed using this model as a starting structure, each domain remained folded and a wide variety of conformations was observed in which the distances between the domains differed markedly (**Supplementary Fig. 6b-g & Supplementary Movies 1-3**). Whilst the three endpoint structures of these simulations satisfy more of the detected inter-domain crosslinks than the crystal structure (an additional 5, **Supplementary Table 3**), they do not satisfy all of the crosslinks observed. However, 18 of the 19 inter-domain crosslinks are compatible with conformations of SurA that were sampled during the three 1 μs simulations (**Supplementary Table 4**), consistent with SurA adopting a broad array of conformations in solution that are in rapid exchange.

We also performed simulated annealing MD simulations of SurA using the detected inter-domain crosslinks as distance restraints (see Methods) in order to visualise possible conformations of the chaperone in which P1 and P2 ate docked onto the core domain (each domain was treated as a rigid body). In the lowest energy structure obtained by this approach (**structure 1** in **Supplementary Fig. 7**) all of the 19 inter-domain crosslinks were satisfied (**Supplementary Fig. 7, Supplementary Table 5**). However, this does not suggest that SurA adopts a unique structure in solution, and indeed other structures obtained in the simulated annealing calculations with different domain orientations explain the observed cross-links almost equally well (the 10 lowest energy structures are shown in **Supplementary Fig. 7, Supplementary Table 5**). In these structures a range of SurA domain orientations are observed (**Supplementary Fig. 7b**), with P2 docking against the core in all 10 structures, whereas P1 adopts a range of conformations (**Supplementary Fig. 7**), consistent with the opening and closing motions observed in smFRET (Fig. 3, **Supplementary Fig. 5**).

Given that this simulated annealing approach will drive SurA to adopt compact states that satisfy the maximum number of restraints within a single structure, more extended states of SurA that are significantly populated in solution, as shown by the smFRET data (Fig. 3), will not be captured by this method. Indeed, as shown by the smFRET and unrestrained MD simulations, the dynamic nature of SurA makes it challenging to define its precise conformational landscape, wherein a broad repertoire of conformations in dynamic exchange on a msec exchange are formed. Notably, no single structure can possibly satisfy the broad distributions observed by smFRET (**Supplementary Fig. 7, Supplementary Table 6**), providing further evidence that the crosslinks observed cannot all result from a single SurA conformation, but result from different rapidly interconverting states. Together, the unrestrained all-atom MD and simulated annealing simulations demonstrate that the three domains of SurA are able to move independently of each other as rigid bodies, facilitated by the flexible linker regions between them (**Supplementary Fig. 1d**). This results in chaperone structures with a broad range of inter-domain distances and orientations.

### SurA binds its OMP substrates at multiple interaction sites

We next investigated how SurA binds its OMP clients, and how this affects conformations adopted by the chaperone. While NMR studies have shown that OMP substrates bound to SurA remain in a dynamic, unfolded state ^40 – 42^, their binding site(s) on SurA remained unexplored. To map the OMP interaction surface on SurA we used *E. coli* OmpX (16 kDa) as a model substrate (OmpX forms an 8-stranded β-barrel in its native state). SurA binds unfolded OmpX with low affinity (K_d,app_ of ~800 nM), as measured by microscale thermophoresis (MST) (**Supplementary Fig. 8**), similar to the affinity of SurA for other OMPs ^28, 43, 44^. SurA-OmpX complexes were assembled by rapid dilution of urea-denatured OmpX into a solution of SurA (final concentrations: 5 μM OmpX, 5 μM SurA, 0.24 M urea) (see Methods) immediately prior to crosslinking with DSBU. A band corresponding to crosslinked SurA-OmpX complexes could be observed by SDS-PAGE (Fig. 4a), and following in-gel digestion a total of 26 unique inter-molecular crosslinked peptides were detected (Fig. 4b, **Supplementary Table 7**). Rather than revealing a unique binding site, the data show that the same residues of OmpX contact different sites on the chaperone surface, consistent with multiple substrate binding modes. Half (13/26) of the crosslinks observed are between OmpX and the SurA core domain (Fig. 4c, **Supplementary Table 7**). Four Lys residues in P2, one in P1, and 7 in the core also crosslinked to OmpX, but no crosslinks were detected for the remaining 11 Lys residues in P1 and P2 indicating a localised binding surface (Fig. 4b,c). Importantly, several crosslinks were detected from *the same residue* in OmpX to *several residues* on SurA (e.g residue 82 of OmpX crosslinks to 13 different residues in SurA spanning all four regions of the chain, **Supplementary Table 7**). Similarly, the *same* site on SurA crosslinked to *multiple sites* on OmpX (e.g. residues 135, 294, 389 and 395 in SurA each crosslink to three residues (50, 71 and 82) in OmpX, **Supplementary Table 7**) consistent with a flexible and dynamic OmpX in the bound state (Fig. 4b). Note that no crosslinks were observed between SurA and Lys112 and Lys122 in OmpX, suggestive of some specificity in the binding interaction, and that since the C-terminal 49 amino acids in OmpX lack Lys residues, no information on whether these regions interact with SurA could be obtained using this crosslinker.

**Fig. 4.**
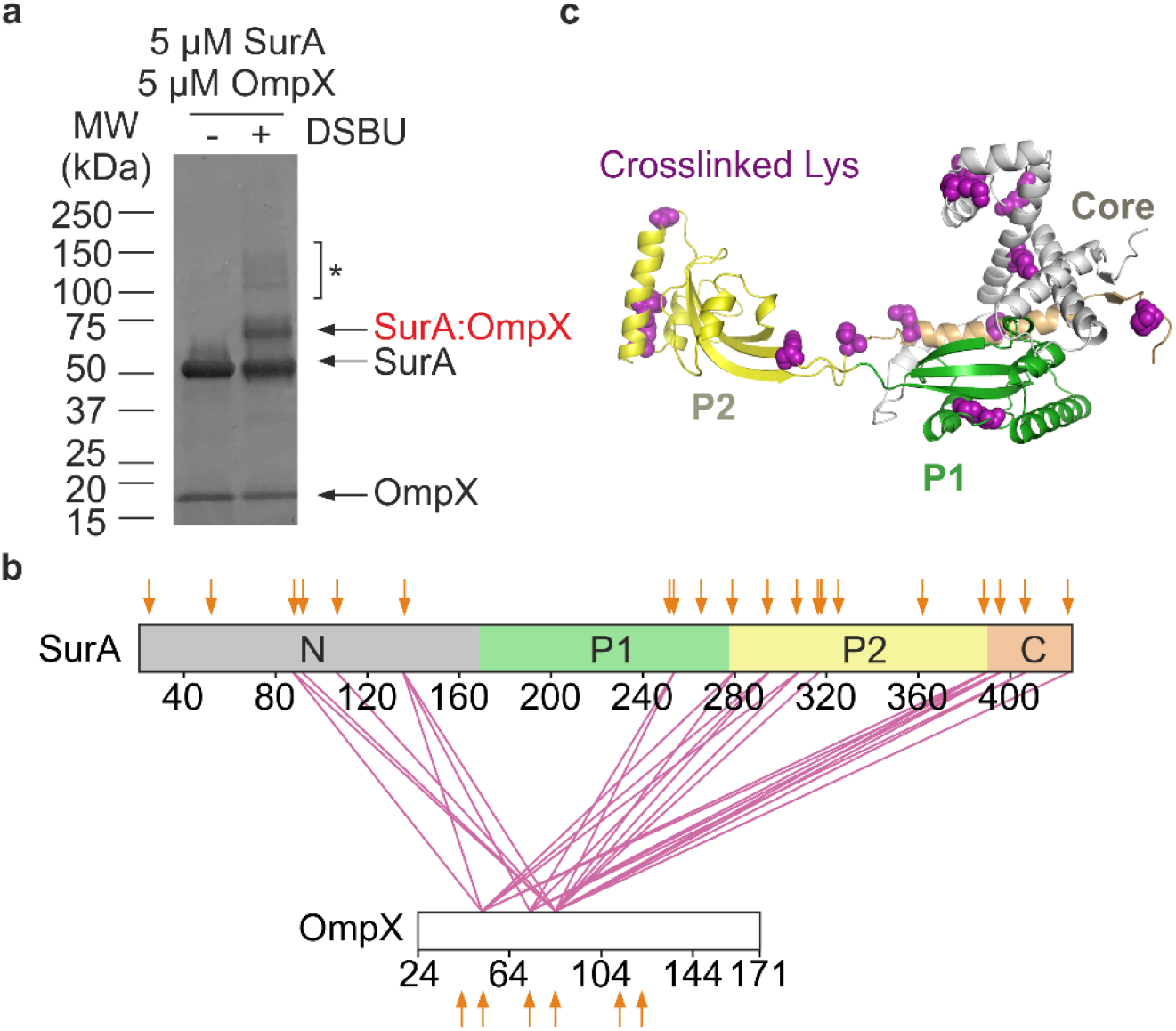
Multi-site binding of OmpX to SurA. **(a)** SDS-PAGE analysis of DSBU crosslinked SurA-OmpX. Note that the species indicated with an asterisk (*) are higher order crosslinked species of mass corresponding to multiple SurA molecules bound to OmpX, consistent with multivalent binding observed previously ^44^. These were not analysed further here. **(b)** Inter-molecular crosslinks detected in the SurA-OmpX complex. The location of all Lys residues are indicated with orange arrows. **(c)** Crystal structure of SurA (PDB 1M5Y ^27^). Purple spheres indicate identified crosslink sites (**Supplementary Table 7**). Samples contained 5 µM SurA, 5 µM OmpX, 0.24 M urea, 50 µM - 2 mM DSBU, in 10 mM potassium phosphate buffer, pH 8.0, 25 °C.

To probe the organisation of the SurA-OmpX complex in more detail we next exploited the ability of the photoactivatable cross-linker MTS-diazirine (**Supplementary Fig. 9a**) to react rapidly (within ns ^45^), and non-specifically with any residue within ~15 Å of the diazirine moiety (Cα-Cα Euclidean distance) ^46^. This “tag transfer” method was developed specifically to enable detection of weak and transient protein-protein interactions ^46^. We created four MTS-diazirine-labelled single-Cys variants of OmpX (M41C, I102C, K122C, V167C), formed complexes of each with SurA, and following rapid UV irradiation (for only 30 sec) ^46^, identified the crosslinked products by LC-MS/MS (Fig. 5a-c, **Supplementary Fig. 9a-c** and **Supplementary Table 8**) ^46^. Despite the location of the labelled Cys residues in distant regions of the OmpX sequence and the lack of specificity of the diazirine crosslinker ^46^, all four Cys-OmpX variants crosslinked to the N-terminal domain of SurA (11 crosslinked sites were identified, **Supplementary Table 8**) and P1 (two crosslinked sites), indicating that these regions form the heart of the binding epitope (Fig. 5b,c, **Supplementary Fig. 9c**). Notably, no crosslinks were detected between OmpX and the SurA P2 domain or C-terminal region, despite the highly promiscuous photoactivatable crosslinker employed. This differs from the SurA-OmpX crosslinks detected with DSBU, probably because a much longer crosslinking time (45 min) required for cross-linking with DSBU. Overall, therefore, the results suggest that OmpX adopts a range of likely interconverting conformations upon binding SurA, in which multiple specific interactions are formed predominantly with the N-terminal region of the chaperone core domain.

**Fig. 5.**
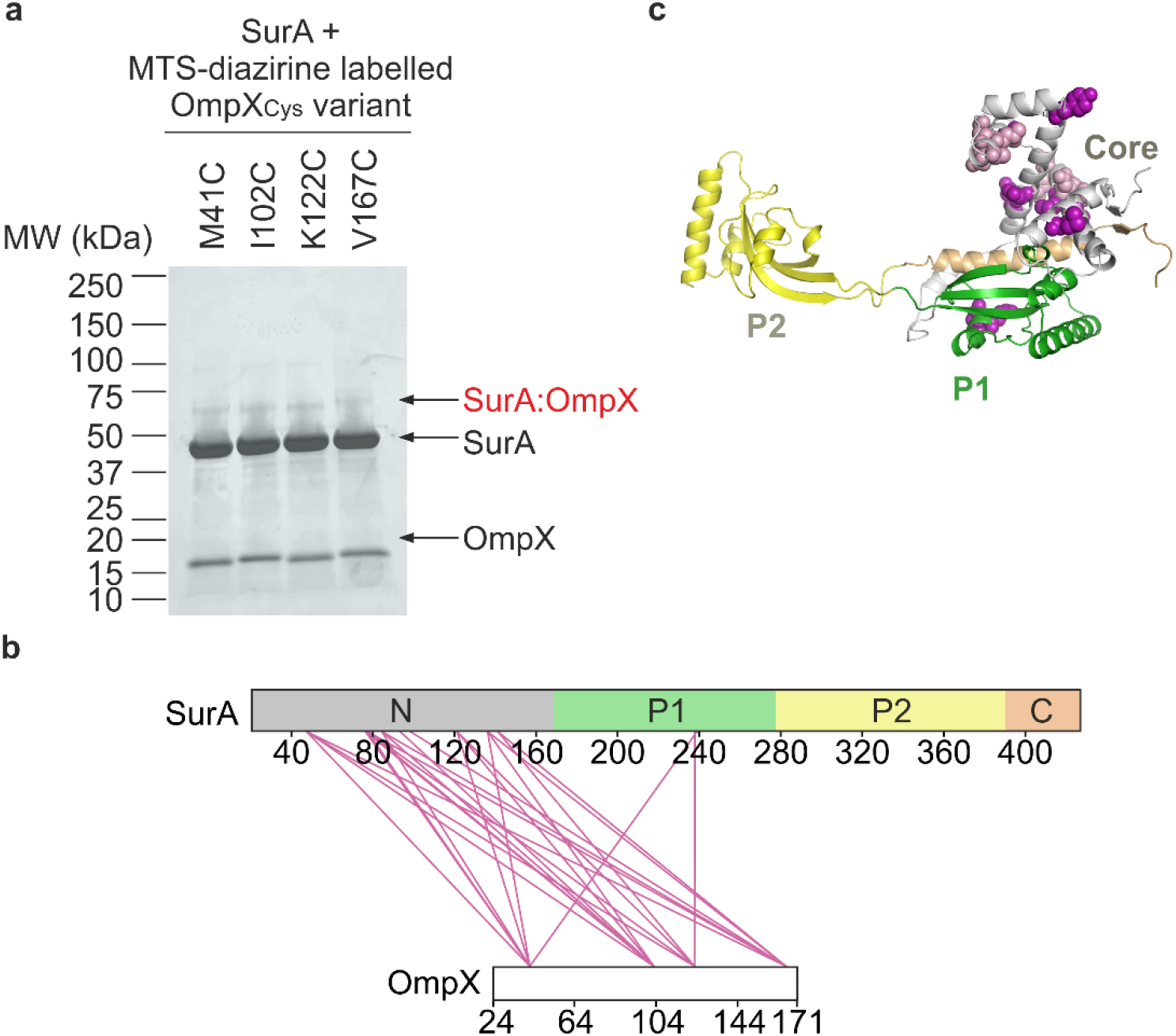
Multiple locations across the OmpX sequence interact with similar sites on SurA. **(a)** Tag-transfer photo-crosslinking ^46^ of SurA-OmpX complexes using OmpX Cys variants labelled with MTS-diazirine analysed by SDS-PAGE. A band corresponding to the SurA-OmpX complex is observed for all OmpX variants following UV irradiation. These bands were not observed when analysed using reducing SDS-PAGE (**Supplementary Fig. 9b**). **(b)** Inter-molecular crosslinks detected in the SurA-OmpX complex. **(c)** Structure of SurA with residues which were photocrosslinked to labelled OmpX Cys variants shown in purple. Where the data quality did not permit residue level assignment, the crosslinked peptide is shown in light purple. Samples contained 10 μM SurA, 5 µM MTS-diazirine-labelled OmpX, 0.24 M urea, in 10 mM potassium phosphate buffer, pH 8.0, 25 °C and crosslinking was initiated by UV LED irradiation of the sample for 30 s (see Methods).

### Conformational changes in SurA upon OMP binding

Next, we examined the conformational changes induced by OMP binding to SurA using differential HDX-MS analysis (Fig. 6a-f and **Supplementary Fig. 10**). We first compared the uptake of deuterium by different regions of SurA in the presence or absence of OmpX under conditions which minimise OMP aggregation (10 mM potassium phosphate, pH 8.0, 0.24 M urea, 4 °C ^28, 47, 48^) (Fig. 6a,b, **Supplementary Fig. 10a**). In the presence of OmpX, regions in SurA that are protected from deuterium uptake upon substrate binding cluster to the core domain. No change in protection in P2 was detected in the presence of OmpX, consistent with the tag transfer XL-MS results and with previous results which have shown that P2 is not required to prevent the aggregation of the small (8-stranded) OmpA ^28^. Intriguingly, two regions of SurA (residues 46-72 in the N-terminal region and 212-239 in P1) (Fig. 6a,b, **Supplementary Fig. 10a**), that are located at the core-P1 interface (**Supplementary Fig. 1d**), were deprotected upon OmpX binding, demonstrating a structural reorganisation of this interface in response to substrate binding.

**Fig. 6.**
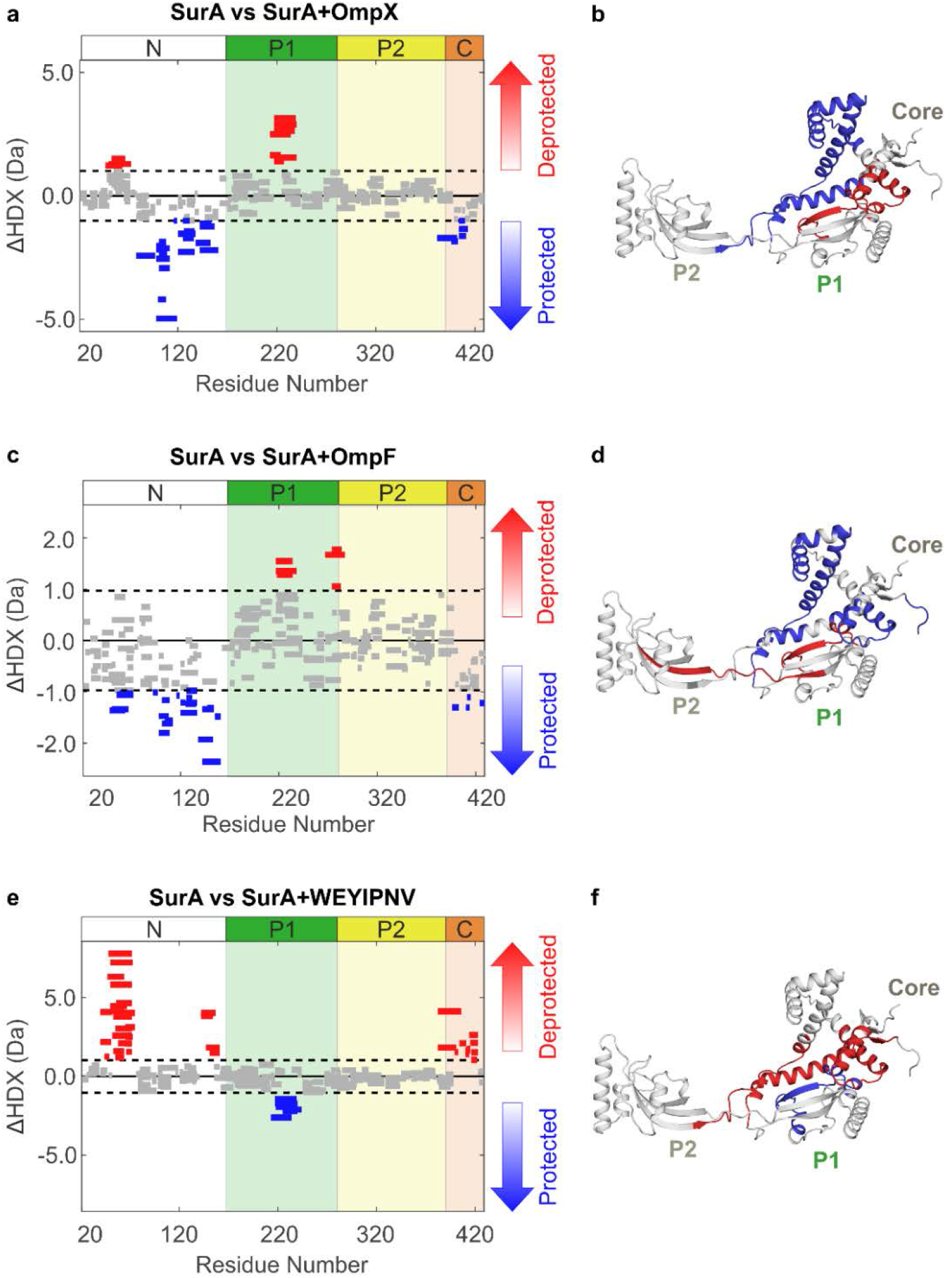
SurA binding to different substrates leads to varying patterns of protection and deprotection by differential HDX-MS analysis. Wood’s plots showing the summed differences in deuterium uptake in SurA over all four HDX timepoints, comparing SurA alone with SurA in the presence of **(a)** OmpX, **(c)** OmpF or **(e)** WEYIPNV. Wood’s plots were generated using Deuteros ^99^. Peptides coloured in blue or red, respectively, are protected or deprotected from exchange in the presence of OmpX/OmpF/WEYIPNV. Peptides with no significant difference between conditions, determined using a 99% confidence interval (dotted line), are shown in grey. Regions of SurA protected or deprotected in the presence of **(b)** OmpX, **(d)** OmpF and **(f)** WEYIPNV coloured in blue or red, respectively. Example deuterium uptake curves are shown in **Supplementary Fig. 10**. See Methods for experimental details.

To determine whether deprotection at the core-P1 interface occurs in the presence of other OMPs, the effects of binding the larger substrate, OmpF (16-stranded), on the HDX properties of SurA was examined. In the presence of OmpF, residues in the N- and C-terminal regions of the core domain were also protected from exchange, consistent with shared OmpX and OmpF binding sites. However, in marked contrast with the results for OmpX in which residues 46-72 of SurA were deprotected from exchange upon substrate binding, these residues were instead protected from exchange in the presence of OmpF, suggesting that the larger OMP binds to, or occludes, a greater surface area on the core (Fig. 6c,d, **Supplementary Fig. 10b**). Importantly, as for OmpX, deprotection was observed in the P1 domain at the core-P1 interface, suggesting that structural reorganisation of this interface also occurs upon OmpF binding. Notably, the hinge region between P1 and P2 (residues 266-286) was also deprotected in the presence of OmpF, suggesting that binding of the larger substrate may also alter the conformational dynamics at locations more distal to the core.

To decouple the phenomena of protection arising as a result of OmpX/OmpF binding and deprotection as a result of conformational changes in SurA, we also compared the levels of deuterium uptake of SurA in the presence of a 7-residue peptide known to bind to the P1 domain (WEYIPNV, K_d_ 1-14 μM) ^49, 50^ (Fig. 6e,f, **Supplementary Fig. 10c,d**). Interestingly, extensive deprotection at the core-P1 interface (residues 39-74, 142-160 and 381-422) was observed in the presence of WEYIPNV, while protection was only observed in P1 at the known peptide binding site (residues 212-243 ^50^) (**Supplementary Fig. 10c,d**). Combined, these results demonstrate that the OMP substrate binding surface is more extensive in OmpX/OmpF compared with WEYIPNV, but for all three cases binding triggers a structural reorganisation between the core and P1 domains.

To further study the effects of substrate binding on the conformations of SurA adopted in solution we used smFRET to examine the inter-domain distances of SurA bound to OmpX, OmpF or WEYIPNV (Fig. 7). Consistent with the HDX data (Fig. 6), binding of OmpX to SurA resulted in changes at the core-P1 interface. Instead of the *ca.* bimodal E_FRET_ distribution observed for the apo-SurA (E_FRET_ centred on ~0.2 and ~0.6, Fig. 3d), a single maximum at an E_FRET_ value (~0.5) between that of the open and closed states was observed for the SurA-OmpX complex (Fig. 7a). By contrast, the E_FRET_ distributions for the core-P2 (Fig. 7b) and P1-P2 (Fig. 7c) SurA variants bound to OmpX were similar to those of apo-SurA. Control experiments in which E_FRET_ was determined for apo-SurA in the presence of 0.24 M urea (used to aid solubilisation of the OMP ^48^) showed only small changes (~10%) in the relative populations for the core-P1_open_ and core-P1_closed_ distributions and increased observation of E_FRET_ values between those of the open and closed states, suggesting more frequent opening/closing transitions in the presence of denaturant. No major differences were observed in the E_FRET_ distributions for the core-P2 and P1-P2 variants upon addition of 0.24 M urea (**Supplementary Fig. 11b,c**). Similar effects on the E_FRET_ distributions were observed when OmpF was added to SurA, (Fig. 7d-f). In marked contrast with the rather modest effects on the EFRET distributions on OmpX/OmpF binding, the addition of the P1-binding peptide WEYIPNV had a profound effect on the core-P1 and core-P2 inter-domain E_FRET_ distributions, inverting the populations of the core-P1_open_ and core-P1_closed_ distributions to favour core-P1_open_ (Fig. 7g), and decreasing the modal E_FRET_ between core-P2 from ~0.34 to ~0.18 (Fig. 7h), without changing the P1-P2 E_FRET_ distribution (Fig. 7i). Consistent with the HDX data, these results suggest that binding of WEYIPNV promotes the release of the P1 domain from the core. BVA on the SurA-substrate complexes indicated that all complexes remained dynamic on the sub-millisecond (sub-ms) timescale (**Supplementary Fig. 12a-i**), although the dynamics of the larger OmpF-SurA complex were dampened relative to those of SurA-OmpX (**Supplementary Fig. 12d-f**).

**Fig. 7.**
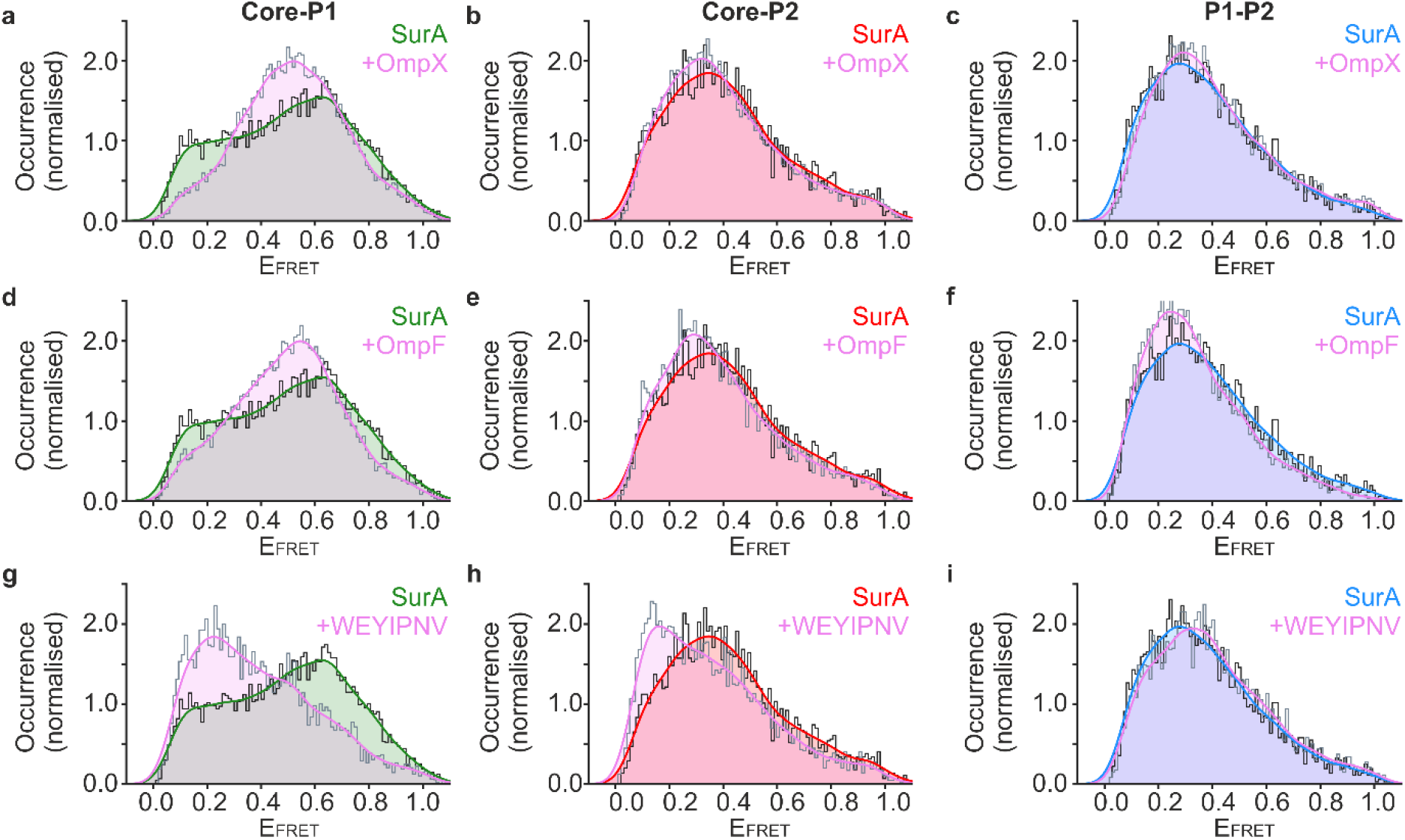
Response of SurA inter-domain distances to substrate binding measured by smFRET. Experimentally measured E_FRET_ distributions at equilibrium for the three pairwise combinations of fluorescently-labelled SurA double mutants (core-P1, core-P2, and P1-P2) in the presence of **(a-c)** OmpX, **(d-f)** OmpF, or **(g-i)** WEYIPNV. Kernel density estimations (KDEs) of the probability density function of the measured E_FRET_ values are shown in green, red and blue for the apo protein core-P1 **(a,d,g)**, core-P2 **(b,e,h)**, and P1-P2 **(c,f,i)** pairwise measurements, respectively, while the corresponding distribution in the presence of OmpX, OmpF or peptide WEYIPNV are shown in violet. Samples contained ~50 pM labelled SurA variant, 1.5 μM OmpX/OmpF/WEYIPNV, in 50 mM Tris-HCl, pH 8.0, 25 °C, with a final urea concentration of 0.24 M in the OMP-containing samples.

## Discussion

Despite its key role in OMP biogenesis and bacterial virulence ^51, 52^, how SurA binds its OMP substrates both specifically, but weakly ^28, 44, 49^, and how it is able to protect its clients from aggregation and deliver them to BAM for folding into the OM, remain poorly understood in molecular detail. Previous NMR studies have shown that OmpX, tOmpA and FhuA are dynamically disordered when bound to SurA ^40 – 42^. However, precisely how SurA binds its OMP clients and how OMP binding alters the conformation(s) adopted by SurA in solution have remained unknown. Here, we have exploited XL, HDX-MS, MD and smFRET, to analyse the conformational dynamics of apo-SurA and to investigate how this is modulated by substrate binding. Further, we have identified the regions of SurA involved in substrate binding for both small (OmpX) and larger (OmpF) clients. The combined data presented are consistent with a model in which specific, yet multi-site, binding by a dynamically disordered substrate is accomplished within a cradle-like conformation of SurA that is very different to that observed in its crystal structure (Fig. 8).

**Fig. 8.**
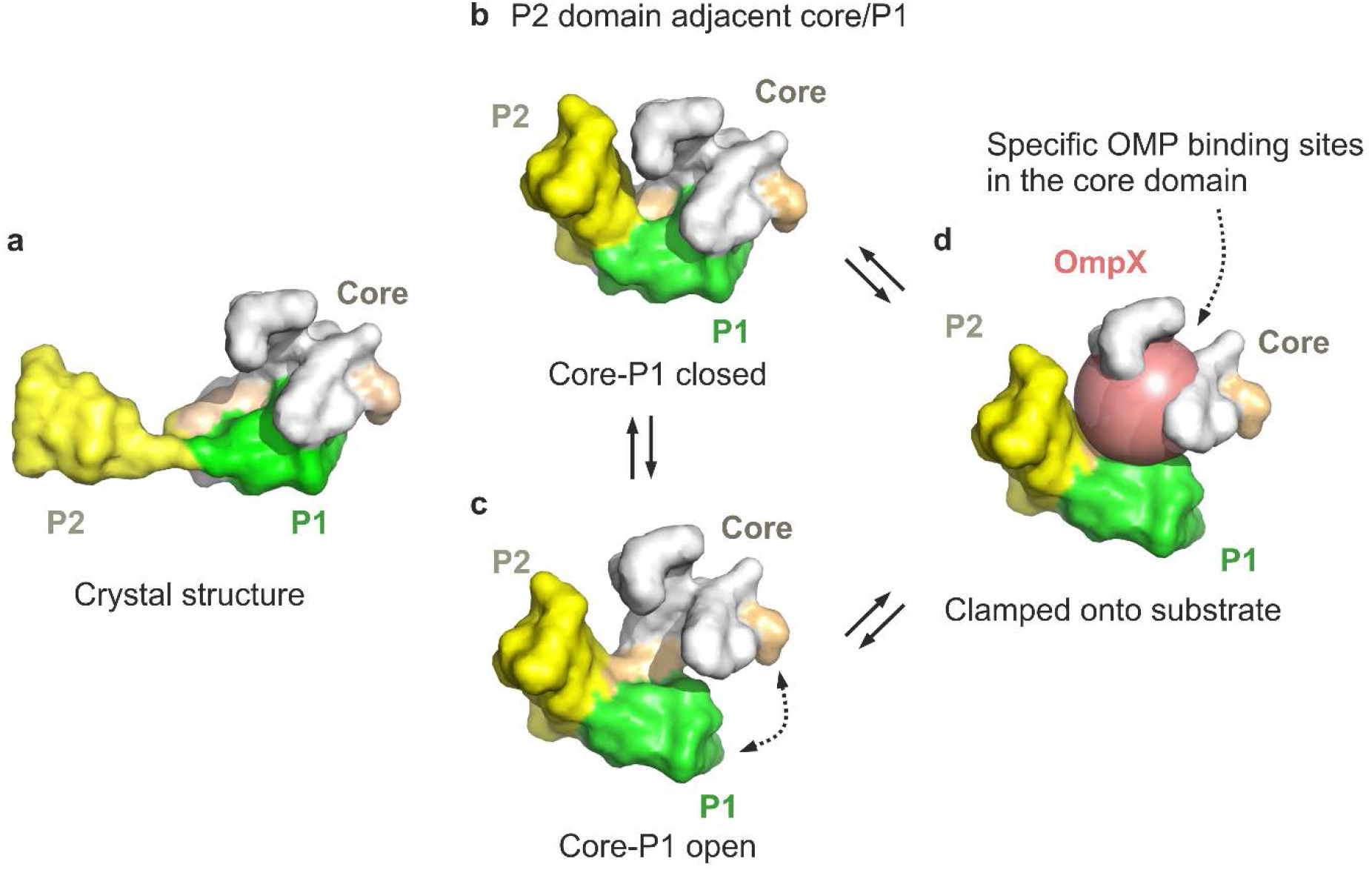
Summary of SurA conformational dynamics and a proposed mechanism of substrate binding. **(a)** The crystal structure of SurA (PDB 1M5Y ^27^). Note that this conformation is not significantly populated in solution, as demonstrated here. Instead, in solution the P2 domain is mostly found close to the core/P1 domains **(b, c)**. In these conformations the P1 domain can adopt **(b)** core-P1_closed_ and **(c)** core-P1_open_ states. **(d)** Substrate binding results in the P1 domain adopting a structure intermediate between core-P1_open_ and core-P1_closed_. Our data are consistent with the OMP client being captured as a dynamically unfolded state ^41^ within a cradle formed by the three domains of SurA. Whether SurA-bound OMP is in a collapsed globule (represented here as a sphere) or a more extended state remains unclear. However, the XL data are consistent with the presence of multiple, specific OMP recognition sites on SurA, suggesting a dynamic ensemble of bound structures. Note that the images presented are schematic and aim to portray the dynamics of the P1, P2 and core domains relative to each other, rather than atomic-level detail.

### SurA: a dynamic ensemble of states primed for OMP capture

Flexibility and/or inter-domain dynamics are key features of the mechanisms of several ATP-independent chaperones, such as Tim9/10 ^53^, Trigger Factor (TF) ^5, 54^, Spy ^9^, SecB ^10^, and the periplasmic OMP chaperones Skp ^55 – 57^ and FkpA ^58^. Like SurA, TF and FkpA are multi-domain proteins containing PPIase domains that exhibit inter-domain dynamics ^58, 59^. The ATP-dependent chaperones GroEL/TriC, Hsp70 and Hsp90 also utilise inter-domain dynamics for their functional cycles ^7, 8, 60^. For the ATP independent chaperones HdeA ^61^ and Hsp33 ^62^, which are activated by acidic and oxidative stress, respectively, conformational switching has been shown to trigger their chaperone function. The results presented here suggest that SurA populates a broad ensemble of structures in solution involving different inter-domain distances and orientations. Interconversion between these conformations occurs on a sub-ms timescale, as demonstrated here using smFRET BVA (**Supplementary Fig. 5**). We propose that such rapid conformational changes are likely to be important for SurA to be able to bind its clients in the periplasm and to release substrates to BAM for folding into the OM. Notably, a similarly broad range of inter-domain motions on comparable timescales to those observed here for SurA was observed previously in MD simulations for the homologous chaperone TF ^5^.

Our combined XL-MS, HDX-MS and smFRET data show that the P1 domain of SurA is not statically bound to the core domain in solution. Instead, these domains are in a dynamic equilibrium between core-P1^open^ and core-P1^closed^ states, suggesting a role for core-P1 dynamics in regulating access of the client OMP to the core chaperone domain and perhaps for access to P1 itself, providing client specificity and enhancing binding affinity ^50^. Previous studies revealed that tethering the core and P1 domains by creation of a disulfide bond impairs OMP assembly *in vivo*, and that destabilising SurA can rescue OMP assembly defects in BAM-compromised strains ^29, 63^. These results can now be explained by the opening and closing motions between the core and P1 domains revealed here by smFRET and HDX-MS. We also show here that the P2 domain commonly populates conformations in close proximity to the core and P1 domains in solution (Figs. 1,2) in marked contrast with the orientation of P2 in the SurA crystal structure in which it is distal to P1/core, most likely as a consequence of crystal packing (**Supplementary Fig. 2**) ^27^. Thus, our data are consistent with SurA frequently adopting a compact, possibly cradle-like, structure (Fig. 8) that may act as the acceptor state for client binding. However, an alternative model in which more extended SurA structures initially capture the unfolded OMP, followed by compaction of the complex, is also possible. In either scenario, the result of binding is a compact SurA in which the client is bound to the core domain, protecting the OMP from aggregation within the dynamic complex. Such a dynamic structure could provide a mechanism for release of bound OMPs to BAM for folding into the OM without the requirement for ATP binding/hydrolysis to drive client release.

### Dynamic interplay between SurA and its substrates

Previous reports based on crystal contacts proposed that SurA may bind its substrates via a binding crevice in the N-terminal domain (Fig. 1) ^27^. By contrast, the results presented here show that SurA binds its OMP clients at multiple sites ^64^, predominantly involving the core domain as indicated by tag-transfer XL and HDX. The additional binding capacity in the P1 and P2 domains may be employed in a substrate-specific manner, as suggested by the finding that P2 is required to suppress the aggregation of the 10-stranded OmpT, but not the 8-strand tOmpA ^28^. Previous results have also shown that deletion of P2 can perturb SurA function *in vivo*, as measured by a decrease in the amount of assembled LamB in a BAM-compromised strain ^29^. These assembly defects are rescued by further deletion of the P1 domain, suggesting that P2 may also play a role in regulating interactions between the P1 and core domains that inhibit SurA chaperone function ^29^. Here we have shown that the three domains of SurA can come into close proximity in solution, consistent with the view that inter-domain communication between all three domains may a play a role in SurA activity. P2 has also been suggested to interact with the BAM complex ^65^, to either localise OMPs to the folding catalyst, prime BAM for OMP insertion, promote substrate release, or create supercomplexes linking SecYEG and BAM across the periplasm ^66^. Differential proteomics experiments have identified reduced levels of eight different OMPs upon SurA deletion ^17^, including OmpX and OmpF which were studied here. Further work will be needed to understand the relay of interactions between SurA, its clients, other chaperones, and BAM, and to discern whether/how the mechanism of OMP delivery to the OM is dependent on the identity of the OMP client.

The XL-MS data presented show that OmpX is able to adopt multiple conformations and orientations when bound to SurA ^40^. The crosslink locations identified on SurA by tag-transfer experiments show no clear correlation with areas with a particular electrostatic potential, hydrophobic patches, or regions of high sequence conservation (**Supplementary Fig. 13**). Instead, they cluster to regions that cluster within the cradle which forms by docking of the three domains of SurA and in which the OMP is sequestered and binds predominantly to the core domain (Fig. 8). Such a model is consistent with *in vivo* data showing that the SurA core domain alone can largely (but not wholly) complement deletion of wild-type SurA ^24, 29, 67^. Specific client interaction sites have also been identified in TF using the substrate PhoA, which are also located in a cradle formed between its domains ^68^. In the presence of substrate, an intermediate state between core-P1_open_ and core-P1_closed_ is observed by smFRET, suggesting that clamp-like motions between these domains may be utilised by SurA to bind and sequester its substrates. This is reminiscent of the lid motions in Hsp70 that entrap its substrates ^69^, the sequestration of OMPs within the cavity of the chaperone Skp ^55, 56^, and the conformational flexibility that has been suggested to be important for substrate capture/release by Spy ^9^.

SurA plays multiple roles in OMP biogenesis, including sequestration of OMPs in the periplasm to prevent their toxic aggregation ^70^ and delivery of OMPs to the BAM complex to enable folding into the OM ^71^. The results presented here demonstrate a role for SurA inter-domain dynamics in OMP binding, notably the reorganisation of the core and P1 domains, and dynamic localisation of P2 close to these domains. Such structural plasticity may also be important for facilitating binding of SurA:OMP complexes to BAM, assisting BAM catalytic activity, or priming the OMP for membrane insertion by pre-selecting favourable conformations for folding, thereby smoothing the energy landscape of folding ^42, 71, 72^. Understanding the interplay between the conformational dynamics of SurA and those of BAM, in particular the communication and coordination between SurA and different BAM subunits, will be essential in unravelling the molecular mechanism of OMP biogenesis. The model of SurA action presented here, whereby a compact, dynamic and responsive chaperone structure is responsible for client binding, represents a first key step in this endeavour. This adds to the growing body of data suggesting that all components of the OMP assembly line, including SurA, Skp ^55, 56^ and BAM ^73 – 77^ have intrinsic conformational dynamics which, in combination, may be key to achieving efficient OMP biogenesis in the absence of ATP.

## Acknowledgements

We thank members of the Radford and Brockwell laboratories for helpful discussions, along with Nasir Khan and James Ault for excellent technical support and Tomas Fessl for advice on FRET data analysis. The plasmid containing the mature sequence of SurA was kindly provided by D. Kahne (Harvard). ANC (BB/P000037/1), BS (BB/N007603/1, BB/T000635/1), MW (BB/N017307/1) and JEH (BB/M011151/1) acknowledge funding from the BBSRC. MW is funded by the EPSRC (EP/N035267/1), JRH was funded by the US National Institutes of Health (GM102829) and PW acknowledges funding from the MRC (MR/P018491/1). The Monolith NT.115 MST instrument was purchased with funding from the Wellcome Trust (105615/Z/14/Z). Funding from the Wellcome Trust (208385/Z/17/Z) and BBSRC (BB/M012573/1) enabled the purchase of mass spectrometry equipment.

## Methods

### Cloning, expression and purification of SurA

A pET28b plasmid containing the mature SurA sequence preceded by an N-terminal 6x His-tag and thrombin-cleavage site (pSK257) was a kind gift from Daniel Kahne (Harvard University, USA) ^78^. The thrombin-cleavage site was mutated to a TEV-cleavage site using Q5 site-directed mutagenesis (NEB), and the resulting plasmid was transformed into BL21(DE3) cells (Stratagene). Cells were grown in LB medium supplemented with 30 µg/mL kanamycin at 37 °C with shaking (200 r.p.m.) until an OD_600_ of ~0.6 was reached. The temperature was subsequently lowered to 20 °C, and expression induced with 0.4 mM IPTG. After ~18 h, cells were harvested by centrifugation, resuspended in 25 mM Tris-HCl, pH 7.2, 150 mM NaCl, 20 mM imidazole, containing EDTA-free protease inhibitor tablets (Roche), and lysed using a cell disrupter (Constant Cell Disruption Systems). The cell debris was removed by centrifugation (20 min, 4 °C, 39,000*g*), and the lysate was applied to 5 mL HisTrap columns (GE Healthcare). The columns were washed with 25 mM Tris-HCl, pH 7.2, 150 mM NaCl and 20 mM imidazole, followed by 25 mM Tris-HCl, 6 M Gdn-HCl, pH 7.2 (to denature the SurA on-column). After washing with 25 mM Tris-HCl, 150 mM NaCl, pH 7.2, SurA was eluted with 25 mM Tris-HCl, 150 mM NaCl, 500 mM imidazole, pH 7.2. The eluate was dialysed against 25 mM Tris-HCl, 150 mM NaCl, pH 8.0 overnight, and the following day TEV protease (purified as described previously ^44^) (*ca.* 0.5 mg) and 0.1 % (v/v) β-mercaptoethanol were added. The cleavage reaction was left to proceed overnight at 4 °C on a tube roller. The cleavage reaction was again applied to the 5-mL HisTrap columns (GE Healthcare) to remove the cleaved His-tag and His-tagged TEV protease. The unbound, cleaved SurA product was dialysed extensively against 25 mM Tris-HCl, 150 mM NaCl, pH 8.0, before being concentrated to ~200 µM with Vivaspin 20 concentrators (Sartorius; 5-kDa MWCO), aliquoted, snap-frozen in liquid nitrogen and stored at −80 °C. Protein concentrations were determined spectrophotometrically using an extinction coefficient at 280 nm of 29450 M^−1^ cm^−1^.

Cys-containing variants (Q85C, N193C, E301C, Q85C-N193C, N193C-E301C and Q85C-E301C) were generated by Q5 site-directed mutagenesis (New England Biolabs) and were purified as detailed above, except for the addition of 10 mM DTT to all buffers in the purification procedure, up until the elution step.

### Cloning of OmpX, OmpF and Cys-OmpX

Codon-optimised synthetic genes (Eurofins) of the mature sequences of OmpX (residues 24-171) and OmpF (residues 23–362) were cloned into pET11a (Novagen) between the NdeI (5′) and BamHI (3′) restriction sites. To create the Cys-OmpX construct, the residues Gly-Ser-Cys were added immediately after the N-terminal Met residue using Q5 site-directed mutagenesis (NEB).

### Expression and purification of OMPs

OMPs were purified using a method adapted from ^48^. Briefly, *E. coli* BL21[DE3] cells (Stratagene, UK) were transformed with a pET11a plasmid containing the gene sequence of the mature OMP. Overnight cultures were subcultured and grown in LB medium (500 ml) supplemented with carbenicillin (100 μg/mL), at 37 °C with shaking (200 r.p.m.). Protein expression was induced with IPTG (1 mM) once an OD_600_ of 0.6 was reached. After 4 h the cells were harvested by centrifugation (5,000 *g*, 15 min, 4°C). The cell pellet was resuspended in 50 mM Tris-HCl pH 8.0, 5 mM EDTA, 1 mM phenylmethylsulfonyl fluoride, 2 mM benzamidine, and the cells were subsequently lysed by sonication. The lysate was centrifuged (25,000 *g*, 30 min, 4°C) and the insoluble material was resuspended in 50 mM Tris-HCl pH 8.0, 2% (*v*/*v*) Triton X-100, before being incubated for 1 h at room temperature, with gentle agitation. The insoluble material was pelleted (25,000 *g*, 30 min, 4°C) and the inclusion bodies washed twice by resuspending in 50 mM Tris-HCl pH 8.0 followed by incubation for 1 h at room temperature with gentle agitation, and then collected by centrifugation (25,000 *g*, 30 min, 4 °C). For the OmpX and OmpF constructs, the inclusion bodies were solubilised in 25 mM Tris-HCl, 6 M Gdn-HCl, pH 8.0 and centrifuged (20,000 *g*, 20 min, 4 °C). The supernatant was filtered (0.2 µM syringe filter, Sartorius, UK) and the protein was purified using a Superdex 75 HiLoad 26/60 gel filtration column (GE Healthcare) equilibrated with 25 mM Tris-HCl, 6 M Gdn-HCl, pH 8.0. Peak fractions were concentrated to ∼500 μM using Vivaspin 20 (5 kDa MWCO) concentrators (Sartorius, UK), and the protein solution was snap-frozen in liquid nitrogen and stored at −80 °C.

### Chemical crosslinking-mass spectrometry (XL-MS)

OmpX was buffer exchanged from storage buffer (25 mM Tris-HCl, 6 M Gdn-HCl, pH 8.0) into 10 mM potassium phosphate, pH 8.0, 8 M urea using Zeba spin desalting columns (Thermo Fisher Scientific, UK). For crosslinking, apo-SurA was prepared at a concentration of 5 µM in 10 mM potassium phosphate, pH 8.0, whilst SurA-OmpX complexes were assembled by mixing SurA and OmpX such that the final concentration of each was 5 µM, in 10 mM potassium phosphate, pH 8.0, 0.24 M urea. DSBU (Thermo Fisher Scientific, UK) was added at 10x (apo-SurA) or 10-400x (SurA:OmpX) molar equivalents relative to the concentration of SurA, and the crosslinking reaction was left to proceed for 45 min at room temperature before quenching by adding 0.2 M Tris-HCl, pH 8.0. The crosslinked material was separated by SDS-PAGE and gel bands corresponding to either SurA alone or the SurA:OmpX complex were excised and the proteins trypsinised in-gel as described previously ^46^. Peptides (5 µL) were injected onto a reverse-phase Acquity M-Class C18, 75 µm x 150 mm column (Waters, UK) and separated by gradient elution of 1-50 % (v/v) solvent B (0.1 % (v/v) formic acid in acetonitrile) in solvent A (0.1 % (v/v) formic acid in water) over 60 min at 300 nL.min^−1^. The eluate was infused into an Orbitrap Q Exactive (Thermo Fisher Scientific, UK) mass spectrometer operating in positive ion mode. Orbitrap calibration was performed using Ultramark solution (Thermo Fisher Scientific, UK). Data acquisition was performed in DDA mode and fragmentation was performed using HCD. Each high-resolution full scan (m/z range 500-2000, R =120,000) was followed by high-resolution product ion scans (R=15,000), with a normalised collision energy of 30 %. The 15 most intense ions in the MS spectrum were selected for MS/MS. Dynamic exclusion of 60 s was used. Crosslink identification was performed using MeroX v1.6.6 ^79^. Raw DSBU XL-MS data are available at the University of Leeds data repository (https://doi.org/10.5518/701).

### Preparation of SurA variants for smFRET

For each SurA variant, the protein was diluted to a concentration of 50 μM in 25 mM Tris-HCl, pH 7.2, 150 mM NaCl, 5 mM DTT. The protein solution was incubated for 30 min at room temperature before being buffer exchanged into 25 mM Tris-HCl, pH 7.2, 150 mM NaCl, 1 mM EDTA using 7 kDa MWCO Zeba spin desalting columns (Thermo Fisher Scientific, UK). A ten-fold molar excess of Alexa Fluor 488 C5 maleimide/Alexa Fluor C5 594 maleimide (Thermo Fisher Scientific, UK) was then added and the samples incubated for 2 h at room temperature with gentle rocking. The reaction was quenched with a 10-fold molar excess (over Alexa Fluor 488 C5 maleimide and Alexa Fluor C5 594 maleimide) of β-mercaptoethanol. Protein was separated from unbound dye by size exclusion chromatography on a Superdex 200 10/300 GL column (GE Healthcare, UK) equilibrated with 50 mM Tris-HCl, 150 mM NaCl, pH 8.0. Fractions containing labelled protein were combined, snap-frozen in liquid nitrogen and stored at −80 °C.

### Single molecule Förster resonance energy transfer (smFRET)

smFRET experiments were performed using a custom-built experimental set-up for μs ALEX as described previously ^80^. Laser wavelengths and powers used were 488 nm, 140 μW and 594 nm, 120 μW, respectively. The laser alternation period was set to 40 μs (duty cycle of 40%). Samples of labelled SurA were prepared on the day of use from concentrated stocks that had been stored at −80 °C and were kept on ice and in the dark while in use. A sample (100 µL, 50 mM Tris-HCl pH 8.0, 50 pM of labelled SurA) selectively supplemented with 1.5 μM OmpX/OmpF and/or 0.24 M urea or 1.5 μM peptide was added atop a coverslip set on the objective. A camera was used to monitor the distance of the focal plane from the coverslip and the objective height adjusted using a piezo-controller (piezo system jena) to 20 μm above the surface of the coverslip. Data acquisition was performed in 3 x 10 min runs with fresh sample prepared after every third collection to counteract the issues of protein aggregation and adherence to the coverslip as well as changes in solution osmolarity resulting from evaporation. Evaporation over the course of 30 mins was minimised by employing a plastic lid that fitted over the coverslip. Data were collected using Labview graphical environment (LabView 7.1 Professional Development System for Windows, National Instruments, Austin, TX) ^81^. Separate photon streams were then converted and stored in an open file format for timestamp-based single-molecule fluorescence experiments (Photon-HDF5), which is compatible with many recent data processing environments ^82^. Fluorescence bursts were analysed using customised Python 2.7 scripts ^83^, and made use of FRETBursts, an open source toolkit for analysis of freely-diffusing single-molecule FRET bursts ^84^. Functions from the FRETBursts package were used to estimate the background signal as a function of time, identify and remove artefacts due to photophysical effects such as blinking, and provide an optimal signal to noise ratio. To obtain EFRET values three correction parameters were applied as described previously ^83^: γ-factor (to account for differences in the efficiency of excitation of each dye), donor leakage into the acceptor channel and acceptor direct excitation by the donor excitation laser. The data from each 10 minute acquisition was merged prior to subsequent analysis. In order to remove bursts arising from incorrectly labelled proteins, the data were filtered using ALEX-2CDE, yielding bursts with a Gaussian distribution of S values in a narrow range of dye stoichiometry (S within 0.25–0.75) ^85^. Typically, ~10000 bursts were collected for each condition examined after all filters had been applied. Filtered bursts were then assembled into 1D histograms and kernel density estimation used to approximate 1D probability density functions of the EFRET values in each condition.

Burst variance analysis (BVA) ^39^ was performed using previously described Python 2.7 scripts ^83^, and plots were made using the Seaborn and Matplotlib ^86^ packages. Visualisations of the available volumes for FRET dyes attached at different positions in SurA were generated using the FRET Positioning and Screening (FPS) software with dye linker lengths and radii parameters suggested in the FPS manual for the FRET dyes used ^37^. Predicted EFRET value distributions from the crystal structure of full-length SurA for each dye pair were calculated from distance distributions generated using the MtsslWizard PyMOL plugin ^87^. Raw smFRET data are available at the University of Leeds data repository (https://doi.org/10.5518/701).

### Fluorescence anisotropy

Fluorescence anisotropy decay measurements were performed on single Cys variants of SurA labelled with Alexa Fluor 488 C5 maleimide or Alexa Fluor C5 594 using a Quantamaster 8000 (Horiba) equipped with a Whitelase supercontinuum pulsed laser (NKT) for excitation with a repetition rate 10 MHz and TCSPC detection. Three pairs of scans were taken with VV and VH polarisation for each sample (500 µL, 600 nM in 50 mM Tris-HCl pH 8), and 10000 photons were collected for each scan. Normalisation and global fitting of each pair of polarised decay curves along with the IRF and HV and HH polarised decays that defined the G factor was performed using FelixGX (Horiba). The steady state and time-resolved anisotropy are related by the following expression:

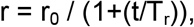

where r = the steady state anisotropy, r_0_ = the initial anisotropy, t = fluorescence lifetime measured for the decay, and T_r_ = the rotational correlation time.

### Molecular dynamics simulations

All-atom molecular dynamics simulations of the mature sequence of SurA (residues 21-428) in explicit solvent were performed with GROMACS 5.0.2 ^88^ using the CHARMM36 force field ^89^. For simulations starting from the crystal structure for full-length SurA (PDB: 1M5Y ^27^), loop residues which are unresolved in the structure were modelled using MODELLER ^90^, and the four missing N-terminal residues were added in Chimera ^91^. The system was minimised (5000 steps) followed by equilibration for 25 ps, with backbone and sidechain position restraints of 400 kJ mol^−1^nm^−2^ and 40 kJ mol^−1^nm^−2^, respectively, in the *x*, *y* and *z* directions. The temperature reached its target value (300 K) within the first 10 ps and remained stable for the rest of the equilibration. The system contained 202 sodium ions and 198 chloride ions (150 mM NaCl), and 70,091 TIP3P water molecules. The total number of atoms was 217,001 in a periodic box size of 13.2 nm x 13.2 nm x 13.2 nm. For simulations starting from a SurA^core-P1-open^ conformation a model was first built using the crystal structures of full-length SurA (PDB: 1M5Y ^27^) and SurA-ΔP2 (PDB: 2PV3 ^50^), in which the P1 domain is extended away from the core. The two structures were aligned on the core domain and the P1 and core domains were removed from the full-length SurA structure. Linker residues between domains were added using MODELLER ^90^, and the four missing N-terminal residues were added in Chimera. The system was minimised (5000 steps) followed by equilibration for 25 ps with backbone and sidechain position restraints of 400 kJ mol^−1^nm^−2^ and 40 kJ mol^−1^nm^−2^, respectively, in the *x*, *y* and *z* directions. The temperature reached its target value (300 K) within the first 10 ps and remained stable for the rest of the equilibration. The system contained 189 sodium ions and 185 chloride ions (150 mM NaCl), and 64,809 TIP3P water molecules. The total number of atoms was 201,129 in a periodic box size of 12.9 nm x 12.9 nm x 12.9 nm. Simulation systems were built using CHARMM-GUI ^92^. In all simulations the pressure was maintained using a Parrinello-Rahman barostat ^93^ and the temperature was maintained using a Nose-Hoover thermostat ^94^. The temperature of the systems was 300 K and the timestep was 2 fs.

Analysis of Cα-Cα distances between residue pairs identified in crosslinking experiments was performed using the ‘*gmx distance*’ GROMACS command. Calculations of solvent accessible surface distances (SASDs) made use of JWalk ^33^. MD simulation data are available at the University of Leeds data repository (https://doi.org/10.5518/701). Included are GROMACS input files, starting structures, reduced MD trajectories and the final structures after 1 μs of simulation.

### Simulated Annealing

Simulated annealing calculations were carried out in XPLOR-NIH ^95^. Crosslinks were treated as distance restraints with a flat-well energy potential using noePot. A rigid-body calculation (100 calculations in total) was performed, where each domain was treated as a rigid body and residues in the linker regions were given torsion angle degrees of freedom. Pseudo-potential energy terms describing covalent geometry restraints were applied to restrict deviation from bond lengths, angles and improper torsion angles. The first step in the structure calculation consisted of 10000 steps of energy minimization, followed by simulated annealing dynamics with all the potential terms active, where the temperature is slowly decreased (3000-25 K) over 4 fs and a final energy minimization in torsion angle space. During the hot phase (T = 3000 K) the crosslink terms were underweighted to allow the domain to sample a large conformational space and they were geometrically increased during the cooling phase. For each calculation the coordinates of P1 and P2 were randomized by applying a random translation within 20 Å and a random rotation within 90° of their initial positions. The linkers were re-built using torsionDB ^96^ to enforce correct geometry before the first step in the structure calculation protocol. Given this simulated annealing approach will drive the structure to compact states, more extended states of SurA that smFRET data show are populated in solution, will not be captured by this method. The structures of the 10 lowest energy conformations of SurA are available at the University of Leeds data repository (https://doi.org/10.5518/701).

### Labelling of Cys-OmpX with Alexa Fluor 488

Purified Cys-OmpX was covalently labelled with Alexa Fluor 488 dye via maleimide chemistry. A sample containing 200 μM Cys-OmpX in 25 mM Tris-HCl, 6 M Gdn-HCl, pH 7.2, was incubated with 10 mM DTT for 30 min. This sample was subsequently buffer-exchanged into 25 mM Tris-HCl, 6 M Gdn-HCl, pH 7.2 (that had been sparged for 15 min with nitrogen gas) using Zeba spin desalting columns (Thermo Fisher Scientific, UK). Alexa Fluor 488 C5 maleimide (Thermo Fisher Scientific, UK) (10 mg/mL dissolved in DMSO) was immediately added to the OmpX sample at a final concentration of 2 mM. The total sample volume was 480 µL. The labelling reaction was kept at 25 °C for 1 h then left overnight at 4 °C. The reaction was then loaded onto a Superdex Peptide 10/300 column equilibrated with 6 M Gdn-HCl, 25 mM Tris-HCl, pH 7.2 to remove the excess free dye. Samples were collected every 1 mL and peak protein fractions tested for dye labelling using a Nanodrop 2000 (Thermo Fisher Scientific, UK). Samples containing labelled OmpX were snap-frozen using liquid nitrogen and stored at −80 °C until required.

### Microscale thermophoresis (MST)

From a 200 μM SurA stock solution in 50 mM Tris-HCl, pH 8.0, a series of two-fold serial dilutions was performed to obtain sixteen 15 µL samples. Labelled Cys-OmpX was buffer exchanged into 8 M urea, 50 mM Tris-HCl, pH 8.0, to a concentration of 1.7 µM. This stock was diluted 16.6-fold to a concentration of 100 nM with 50 mM Tris-HCl, pH 8.0, then immediately added to the sixteen SurA-containing samples in 15 µL aliquots (30 µL total sample volume). The final sample concentrations were 50 nM Cys-OmpX, 100 µM – 3 nM SurA, 0.24 M urea, 50 mM Tris-HCl, pH 8.0. Samples were immediately added to capillaries by capillary action then read using a Monolith NT.115 MST instrument (NanoTemper, Germany). To obtain the dissociation constant, K_d_, data were fitted to the Hill equation:

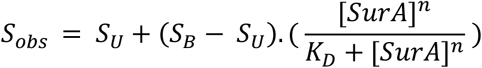

where S_obs_ is the observed signal, S_U_ is the signal from unbound OmpX, S_B_ is the signal from bound OmpX, and n is the Hill coefficient. Data fitting was carried out using IgorPro 6.3.4.1 (Wavemetrics, Oregon, USA).

### Tag-transfer photocrosslinking

Single Cys variants of OmpX (M41C, I102C, K122C, V167C) were conjugated with MTS-diazirine, as described previously ^46^. SurA-OmpX complexes were assembled by mixing SurA with each OmpX variant such that the final concentrations of each was 10 µM and 5 µM, respectively, in 10 mM potassium phosphate buffer, pH 8.0, 0.24 M urea. Photocrosslinking was performed for 30 sec using a UV LED irradiation platform ^46^. The crosslinked material was separated by SDS-PAGE. The gel band corresponding to the crosslinked complex was excised and the proteins were trypsinised in-gel, as described previously ^46^. To detect additional modified peptides, reduction of the crosslinker and thiol capture was performed to enrich crosslinked peptides. Peptides (5 µL) were injected onto a reverse-phase Acquity M-Class C18, 75 µm x 150 mm column (Waters, UK) and separated by gradient elution of 1-50 % (v/v) solvent B (0.1 % (v/v) formic acid in acetonitrile) in solvent A (0.1 % (v/v) formic acid in water) over 60 min at 300 nL.min^−1^. The eluate was infused into a Xevo G2-XS (Waters, UK) mass spectrometer operating in positive ion mode. Mass calibration was performed by infusion of aqueous NaI (2 µg/µL). [Glu1]-Fibrinopeptide B (GluFib) was used for the lock mass spray, with a 0.5 s lock spray scan taken every 30 s. The lock mass correction factor was determined by averaging 10 scans. Data acquisition was performed in DDA mode with a 1 s MS scan over m/z 350-2000. The 4 most intense ions in the MS spectrum were selected for MS/MS by CID, each with a 0.5 s scan over m/z 50-2000. The collision energy applied was dependent upon the charge and mass of the selected ion. Dynamic exclusion of 60 s was used. Data processing and modification localization was performed using PEAKS Studio 8.5 (Bioinformatics Solutions). Raw tag-transfer MS data are available at the University of Leeds data repository (https://doi.org/10.5518/701).

### Hydrogen-deuterium exchange mass spectrometry

An automated HDX robot (LEAP Technologies, Ft Lauderdale, FL, USA) coupled to a Acquity M-Class LC and HDX manager (Waters, UK) was used for all HDX-MS experiments. For differential HDX-MS of SurA in the absence and presence of OmpX/OmpF, samples contained 8 µM of SurA or 8 µM of SurA with 8 µM OmpX/OmpF (in 10 mM potassium phosphate, pH 8.0, 0.24 M urea). For differential experiments with addition of WEYIPNV peptide, the samples contained 8 µM SurA with 110 µM peptide (in 10 mM potassium phosphate, pH 8.0). Note that the addition of 0.24 M urea does not dramatically alter the intrinsic rate of exchange ^97^.

30 μL of protein-containing solution was added to 135 μL deuterated buffer (10 mM potassium phosphate buffer pD 8.0, 0.24 M d_4_-urea or 10 mM potassium phosphate buffer pD 8.0, 82 % D_2_O) and incubated at 4 °C for 0.5, 2, 30 or 120 min. Four replicate measurements were performed for each condition and each time point. After labelling, HDX was quenched by adding 100 μL of quench buffer (10 mM potassium phosphate, 2 M Gdn-HCl, pH 2.2) to 50 μL of the labelling reaction. 50 μL of the quenched sample was passed through immobilised pepsin and aspergillopepsin columns (Affipro, Mratín, Czech Republic) connected in series (20 °C) and the peptides were trapped on a VanGuard Pre-column [Acquity UPLC BEH C18 (1.7 μm, 2.1 mm × 5 mm, Waters, UK)] for 3 min. The peptides were separated using a C18 column (75 μm × 150 mm, Waters, UK) by gradient elution of 0–40% (v/v) acetonitrile (0.1% v/v formic acid) in H_2_O (0.3% v/v formic acid) over 7 min at 40 μL min^−1^. Peptides were detected using a Synapt G2Si mass spectrometer (Waters, UK). The mass spectrometer was operated in HDMS^E^ mode, with dynamic range extension enabled (data independent analysis (DIA) coupled with IMS separation) were used to separate peptides prior to CID fragmentation in the transfer cell ^98^. CID data were used for peptide identification, and uptake quantification was performed at the peptide level (as CID results in deuterium scrambling). Data were analysed using PLGS (v3.0.2) and DynamX (v3.0.0) software (Waters, UK). Restrictions for peptides in DynamX were as follows: minimum intensity = 1000, minimum products per amino acid = 0.3, max sequence length = 25, max ppm error = 5, file threshold = 3. The software Deuteros ^99^ was used to identify peptides with statistically significant increases/decreases in deuterium uptake (applying a 99 % confidence interval) and to prepare Wood’s plots. Raw HDX-MS data are available at the University of Leeds data repository (https://doi.org/10.5518/701).

### Analysis of electrostatic surface potential and residue conservation of SurA

Calculation of the surface electrostatic potential of SurA was performed using the APBS plugin for PyMOL ^100^. Amino acid conservation analysis was carried out using the ConSurf webserver using default parameters ^101^.

