## Supplementary Information for "Inter-domain dynamics in the chaperone SurA and multi-site binding to its unfolded outer membrane protein clients"

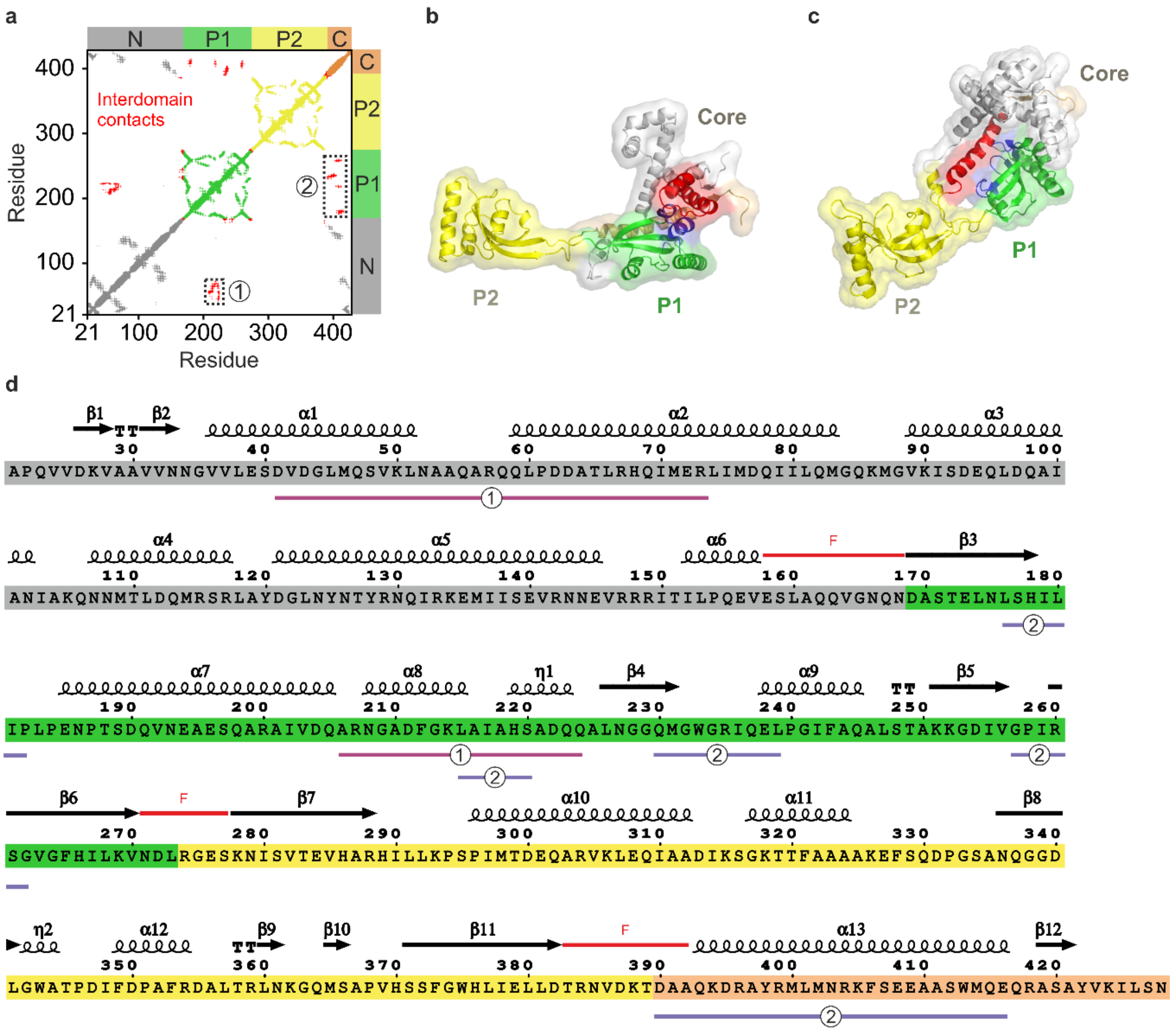


**Supplementary Fig. 1. Inter-domain contacts and secondary structure of *E. coli* SurA. (a)** Contact map showing residue pairs with a Cα-Cα distance of less than 12 Å. Contacts within the core (N- and C-terminal regions), P1 and P2 domains are coloured grey, green and yellow, respectively. Inter-domain contacts are shown in red. Contacts between the core and P1 domains cluster in two regions (shown by boxed regions 1 and 2). These clusters correspond to contacts between P1 and the N-terminal region of the core domain (cluster 1), and contacts between P1 and the C-terminal helix (cluster 2). **(b)** Inter-domain contacts from cluster 1 in **(a)**, involving contacts between P1 and the N-terminal region of the core domain. Residues involved in inter-domain contacts in the core and P1 domains are coloured in red and blue, respectively. **(c)** Inter-domain contacts from cluster 2 in **(a)**, summarising contacts between P1 and the C-terminal helix of the core domain. Residues are coloured as in **(b)**. **(d)** SurA sequence coloured by domains (grey: N-terminal region of the core domain, green: P1, yellow: P2, orange: C-terminal region of the core domain). Regions of secondary structure are shown above the sequence: α-helices (α), β-strands (β), $3_{10}$-helices (η) and turns (T). Flexible regions (F) linking domains are indicated with red lines and the two clusters (1 and

2) of contacts from **(a-c)** are indicated. The figure was prepared using the ENDscript server [1].

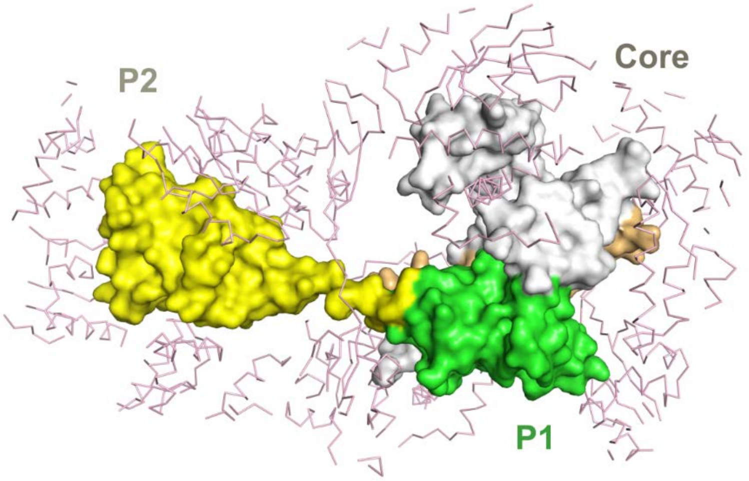


**Supplementary Fig. 2. Multiple packing interactions in the crystal structure of full-length SurA (PDB: 1M5Y [2])**. One copy of full-length SurA is shown in surface representation with the N-terminal region of the core, P1, P2, and the C-terminal region of the core shown in grey, green, yellow and orange, respectively. Atoms from neighbouring molecules in the crystal within 20 Å are shown in pink in ribbon representation.

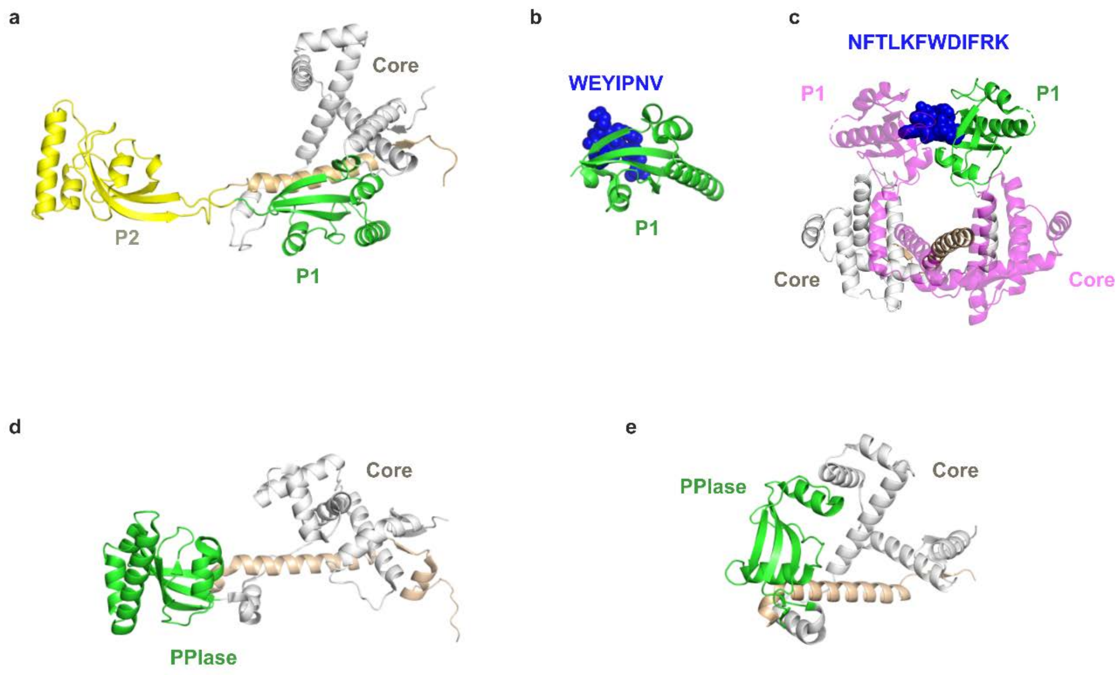


**Supplementary Fig. 3. Crystal structures of *E. coli* SurA, SurA-peptide complexes and SurA homologues. (a)** Structure of *E. coli* SurA (PDB: 1M5Y [2]). **(b)** Structure of the P1 domain of *E. coli* SurA bound to the peptide WEYIPNV (blue spheres) (PDB: 2PV1 [3]). **(c)** Structure of dimeric *E. coli* SurA-ΔP2 in complex with the peptide NFTLKFWDIFRK (blue spheres) (PDB: 2PV3 [3]). For clarity, one monomer is coloured as in **(a)** and the other is shown in pink. **(d)** Structure of the SurA homologue LIC12922 from *Leptospira interrogans (*PDB: 3NRK) [4]. **(e)** Structure of SurA homologue Cj1289 from *Campylobacter jejuni*, (PDB: 3RGC) [5]. Note that *E. coli* SurA homologues from γ- and β- proteobacteria commonly contain two PPIase domains, whilst those from α- ε- and δ-proteobacteria more commonly contain only one PPIase domain [6]. *C. jejuni* is a member of the ε-proteobacterial class, and *Leptospira interrogans* is a member of the phylum Spirochaetes.

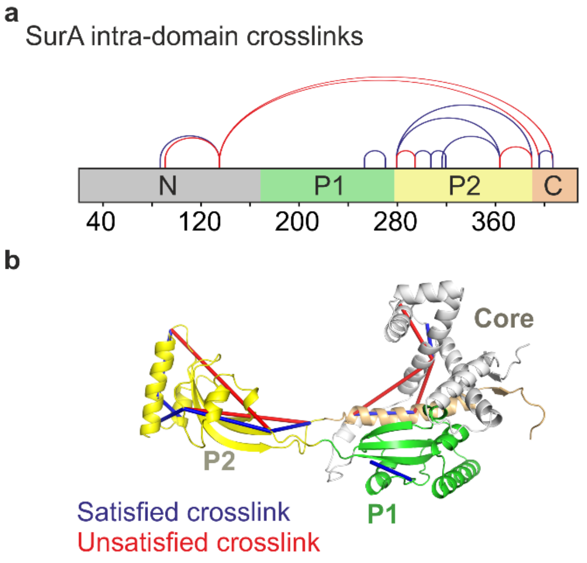


**Supplementary Fig. 4. Intra-domain Lys-Lys cross-links in apo-SurA. (a)** Locations of the 13 experimentally observed SurA intra-domain crosslinks. Crosslinks between residues with a solvent accessible surface distance (SASD) of less than or greater than 35 Å in the crystal structure of full-length SurA (PDB: 1M5Y [2]) are defined as satisfied (blue) (8 crosslinks) or violated (red) (5 crosslinks), respectively. **(b)** Locations of all intra-domain cross-links shown on the SurA crystal structure (PDB 1M5Y [2]). Note that for clarity crosslinks are shown as straight lines between residues, rather than solvent accessible distance paths, coloured as in **(a)**. Details of crosslinked residues are given in **Supplementary Table 1**.

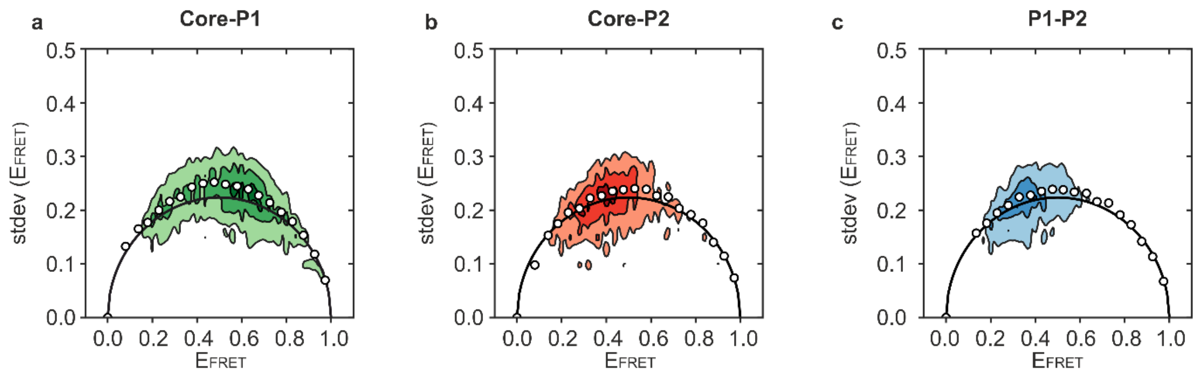


**Supplementary Fig. 5. Burst variance analysis (BVA) of smFRET data for apo-SurA indicates sub-millisecond inter-domain dynamics.** BVA analysis for apo-SurA for all three pairwise combinations of fluorescently-labelled SurA double mutants, **(a)** core-P1, **(b)** core-P2, and **(c)** P1-P2. Each detected burst was binned into sub-bursts containing 5 photons. $E_{FRET}$ values were calculated for each sub-burst and the $E_{FRET}$ value for each burst plotted against the standard deviation of the sub-burst $E_{FRET}$ values within the burst. The black lines indicate the expected shot-noise limited standard deviation as a function of $E_{FRET}$. The average values of the measured variance (white circles) above the theoretical distribution indicate dynamics on a timescale faster than the duration of the bursts (here sub-ms).

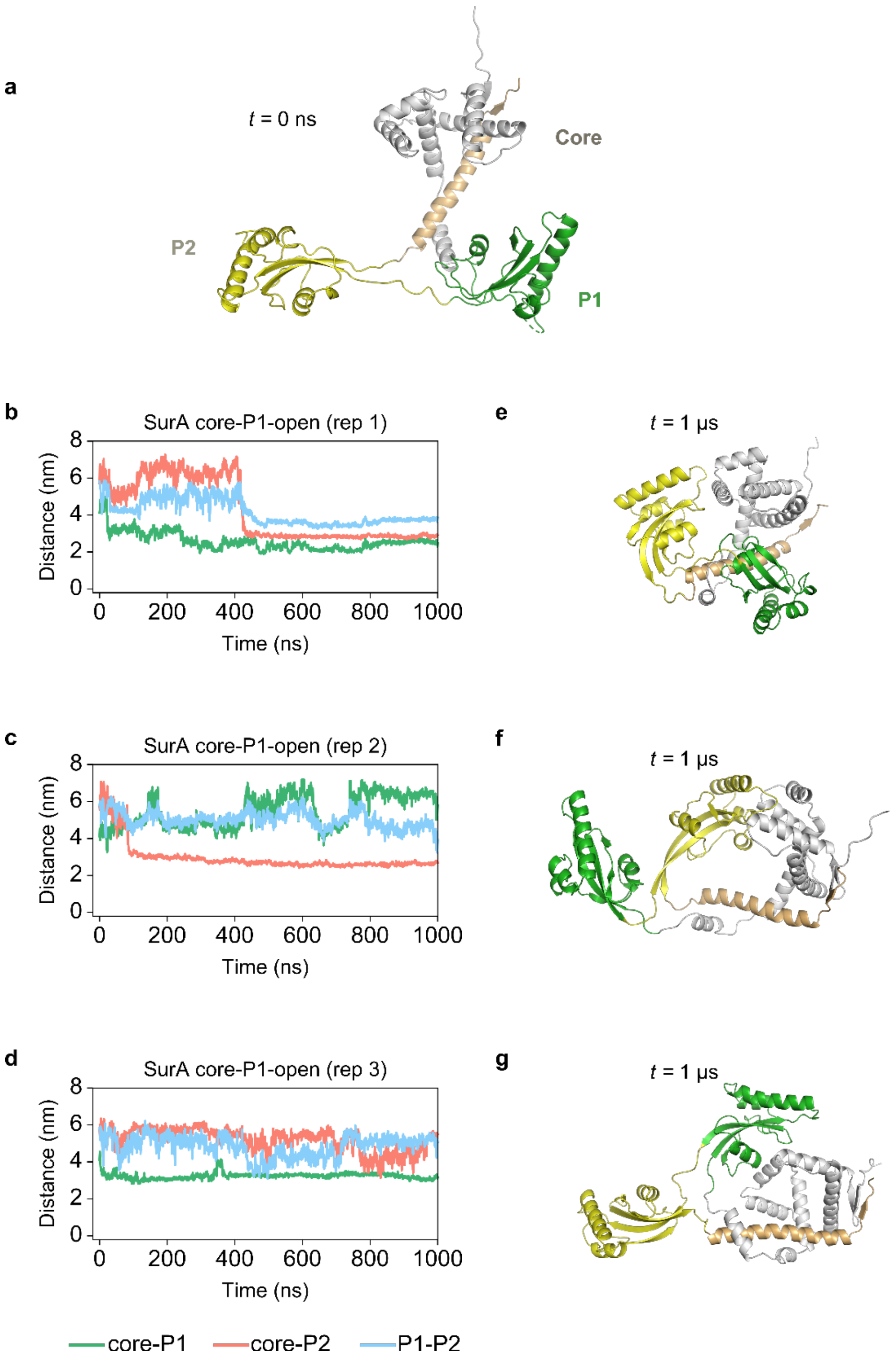


**Supplementary Fig. 6. MD simulations of SurA$^{\text{core-P1-open}}$ suggest a broad conformational ensemble with a range of inter-domain orientations and distances. (a)** Model of a SurA$^{\text{core-P1-open}}$ conformation used as a starting structure for simulation. The model was generated using the X-ray structures of full-length SurA (PDB: 1M5Y [2]) and SurA-ΔP2 (PDB: 2PV3 [3]) (see Methods). **(b-d)** Inter-domain distances over time for each of 3 x 1 μs simulations starting from the SurA$^{\text{core-P1-open}}$ model shown in **(a)**. Inter-domain distances between the centre of mass of each domain for core-P1, core-P2 and P1-P2, are shown in green, red and blue, respectively. **(e-g)** Structures at the end of 1 μs simulation for each of replicates 1, 2 and 3, respectively. Rep: repeat simulation.

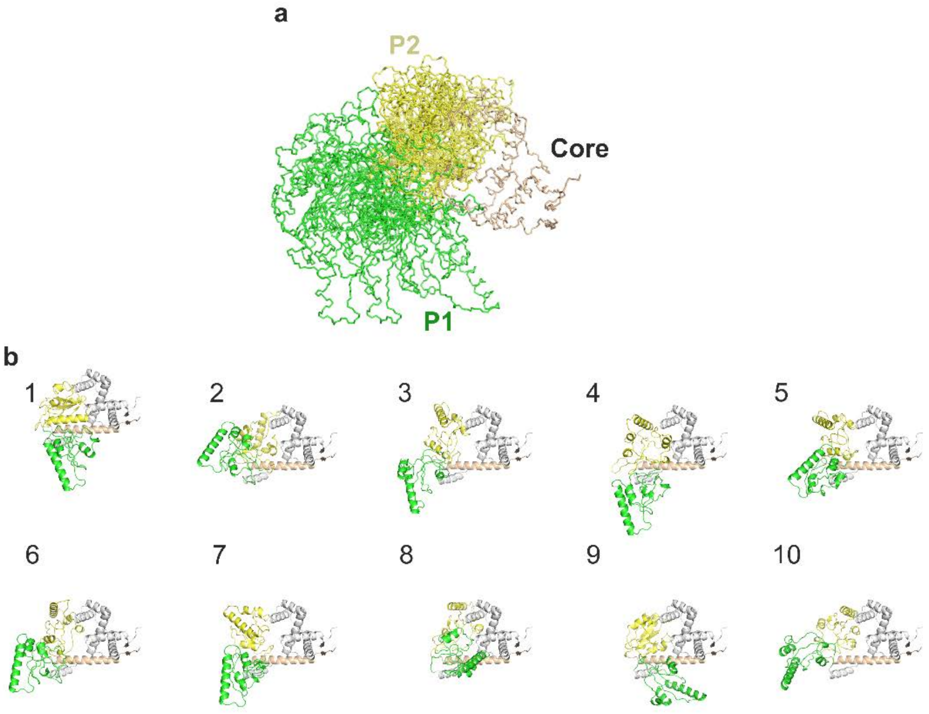


**Supplementary Fig. 7. Simulated annealing MD results in structures consistent with the XL-MS and smFRET data. (a)** Overlay of the 10 lowest energy structures of SurA from simulated annealing MD. The structures are all aligned on the core domain. **(b)** The 10 lowest energy structures of SurA from simulated annealing MD.

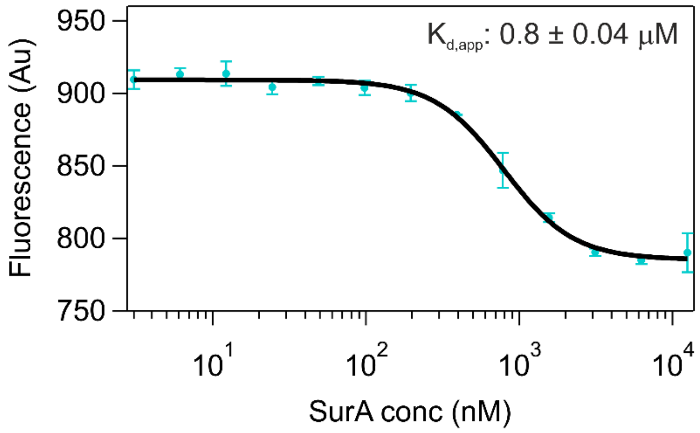


**Supplementary Fig. 8. OmpX binds to SurA with μM affinity.** Microscale thermophoresis (MST) data for binding of SurA to OmpX. Samples contained 50 nM Alexa Fluor 488-labelled OmpX (see Methods), SurA (0.3 nM-12.5 μM), 0.24 M urea, 50 mM Tris-HCl, pH 8.0, 25 °C. A fit to the Hill equation is indicated by a black solid line. Data are shown as the mean ± standard deviation of three independent experiments. The fitted values for $K_{d,app}$ and Hill coefficient were 0.8 ± 0.04 μM and 1.5 ± 0.1, respectively.

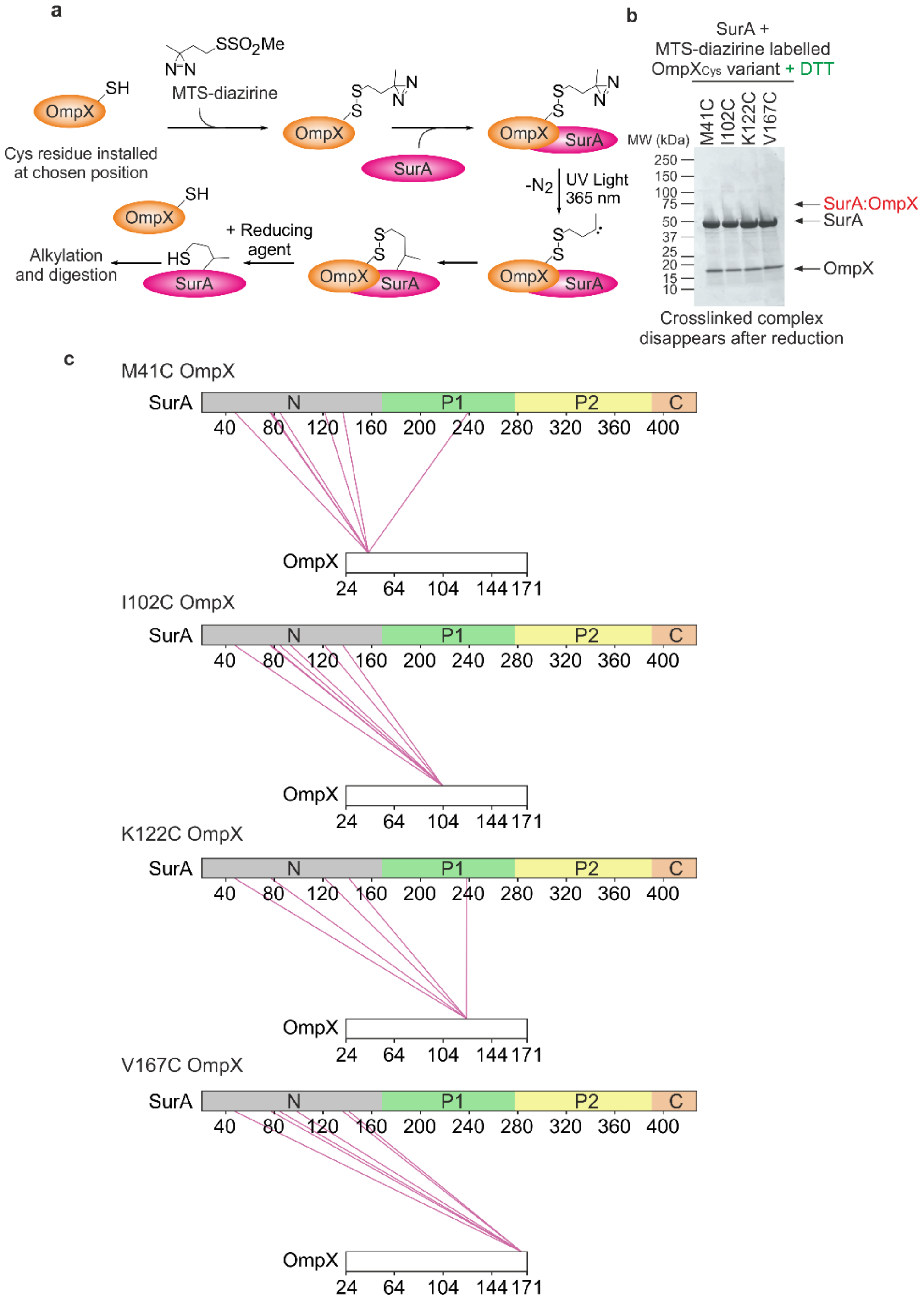


**Supplementary Fig. 9. Multiple locations across the OmpX sequence interact with similar sites on SurA. (a)** Outline of the ‘tag-transfer’ photo-crosslinking workflow (see Methods) [7]. **(b)** SDS-PAGE analysis of photo-crosslinking reactions between SurA and ‘tagged’ OmpX variants under reducing conditions. Note that the band corresponding to the SurA-OmpX complex is lost in the presence of reducing agent (compare with **Fig. 5a**). **(c)** Crosslinks identified for each of the Cys residues introduced into the OmpX sequence. Note that the combined dataset for all four Cys variants are shown in **Fig. 5c**.

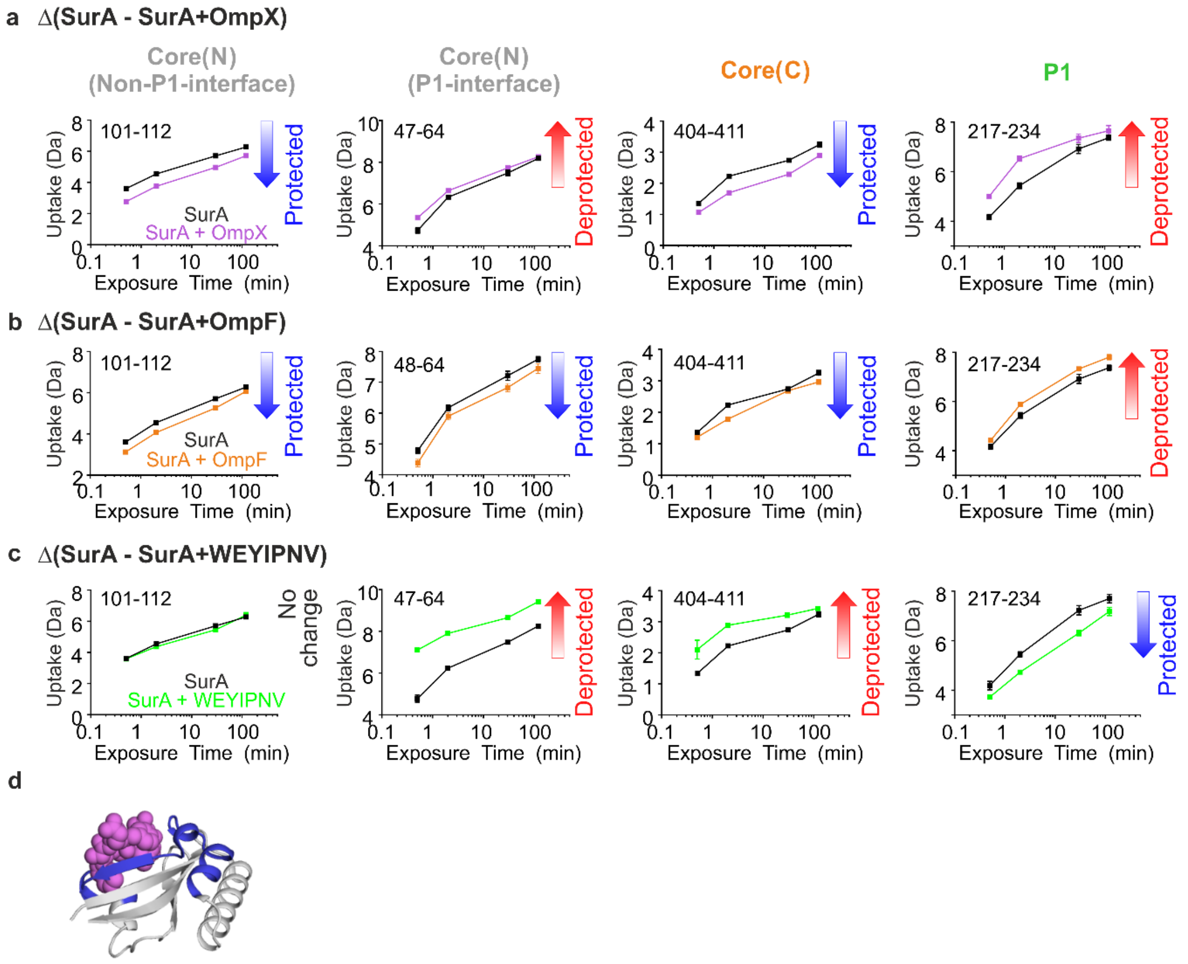


**Supplementary Fig. 10. Different regions of SurA are protected or deprotected from hydrogen exchange in the presence of OmpX, OmpF or WEYIPNV.** Example deuterium uptake curves for SurA (black) or SurA in the presence of **(a)** OmpX, **(b)** OmpF or **(c)** WEYIPNV (coloured as indicated) for four regions of SurA: (left) N-terminal region of the core domain distal to the core-P1 interface, (second from left) N-terminal region of the core domain at the core-P1 interface, (third from left) the C-terminal region of the core domain, and (right), the P1 domain at the interface with the core. The residue numbers in each peptide are indicated in the top left of each plot. **(d)** Crystal structure of WEYIPNV bound to the P1 domain of SurA (PDB 2PV1 [3]). The peptide is shown as purple spheres, and the region of SurA protected from HDX in the presence of WEYIPNV is shown in blue. See Methods for experimental details.

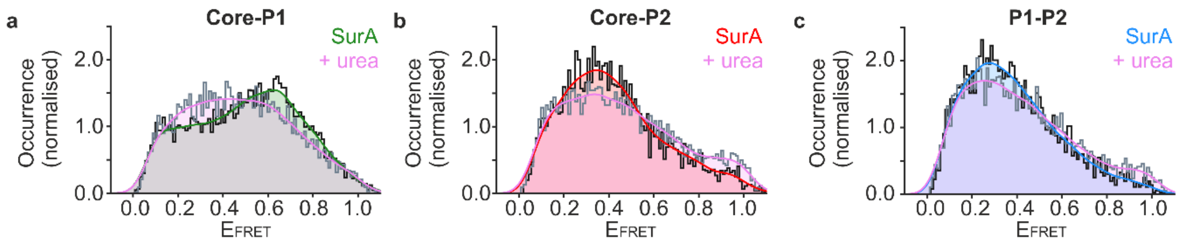


**Supplementary Fig. 11. Addition of 0.24 M urea to SurA alters the equilibrium between core-P1$_{open}$ and core-P1$_{closed}$** conformations. Experimentally measured $E_{FRET}$ distributions for the three pairwise combinations of fluorescently-labelled SurA double mutants (core-P1, core-P2, and P1-P2) in the presence of 0.24 M urea are shown in **(a)**, **(b)**, and **(c)**, respectively. In each panel, kernel density estimations (KDEs) of the probability density function of the measured $E_{FRET}$ values are shown in pink (for data acquired in the presence of 0.24 M urea). For comparison, KDEs in the absence of urea (**Fig. 3**) are overlaid and are shown in green, red and blue for **(a)** core-P1, **(b)** core-P2, and **(c)** P1-P2, respectively. Samples contained ~50 pM labelled SurA variant in 0.24 M urea, 50 mM Tris-HCl, pH 8.0, 25 ºC.

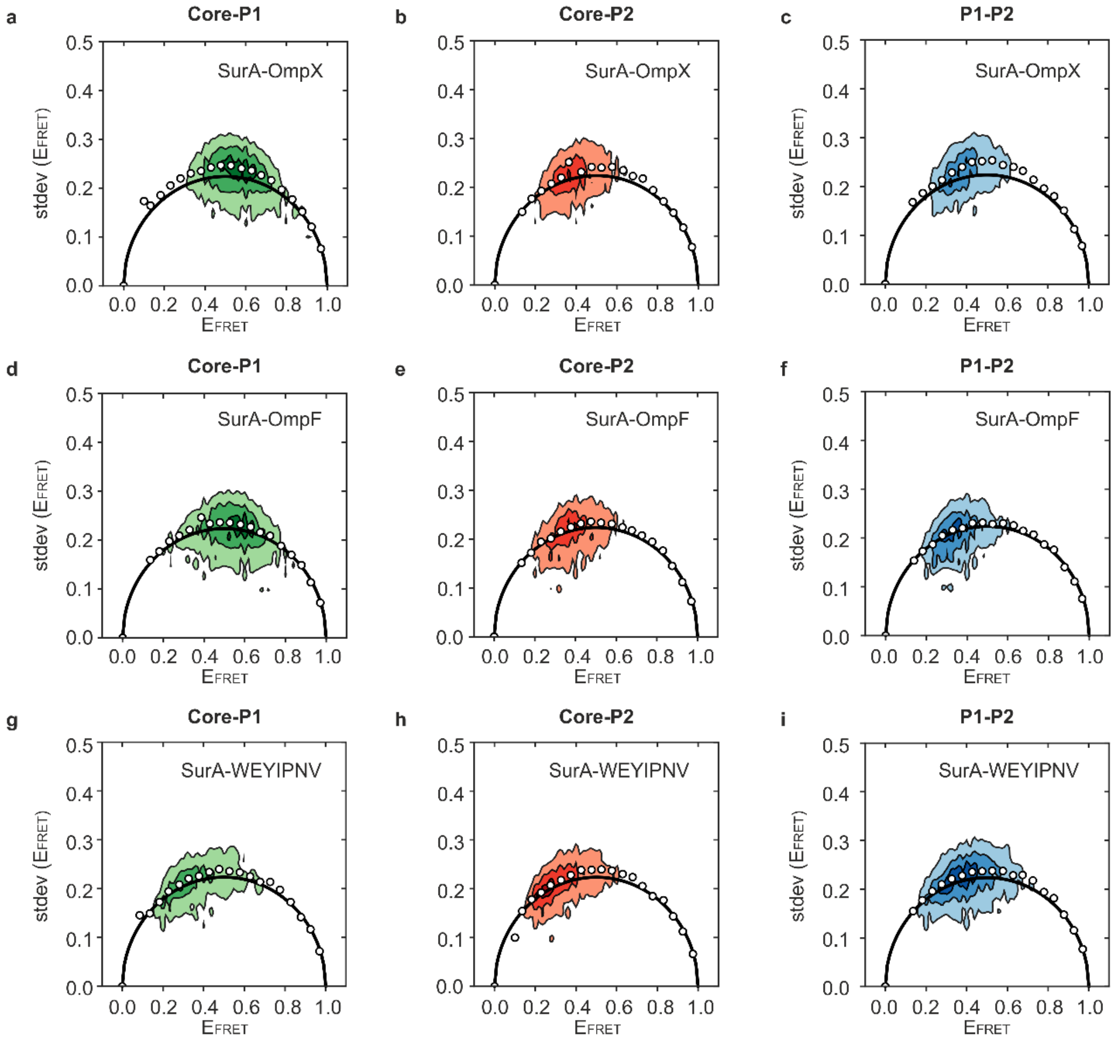


**Supplementary Fig. 12. Burst variance analysis of smFRET data for SurA in complex with OmpX, OmpF or the peptide WEYIPNV indicates sub-millisecond inter-domain dynamics.** BVA analysis for SurA in the presence of **(a-c)** OmpX, **(d-f)** OmpF and **(g-i)** WEYIPNV for all three pairwise combinations of fluorescently-labelled SurA double mutants **(a,d,g)** core-P1, **(b,e,h)** core-P2, or **(c,f,i)** P1-P2. Each detected burst was divided into sub-bursts each containing 5 photons. $E_{FRET}$ values were calculated for each sub-burst and the $E_{FRET}$ value for each burst plotted against the standard deviation of the sub-burst $E_{FRET}$ values within the burst. The black lines indicate the expected shot-noise limited standard deviation as a function of $E_{FRET}$. The average values of the measured variance (white circles) above the theoretical distribution indicate dynamics on a timescale faster than the duration of the bursts (here sub-ms).

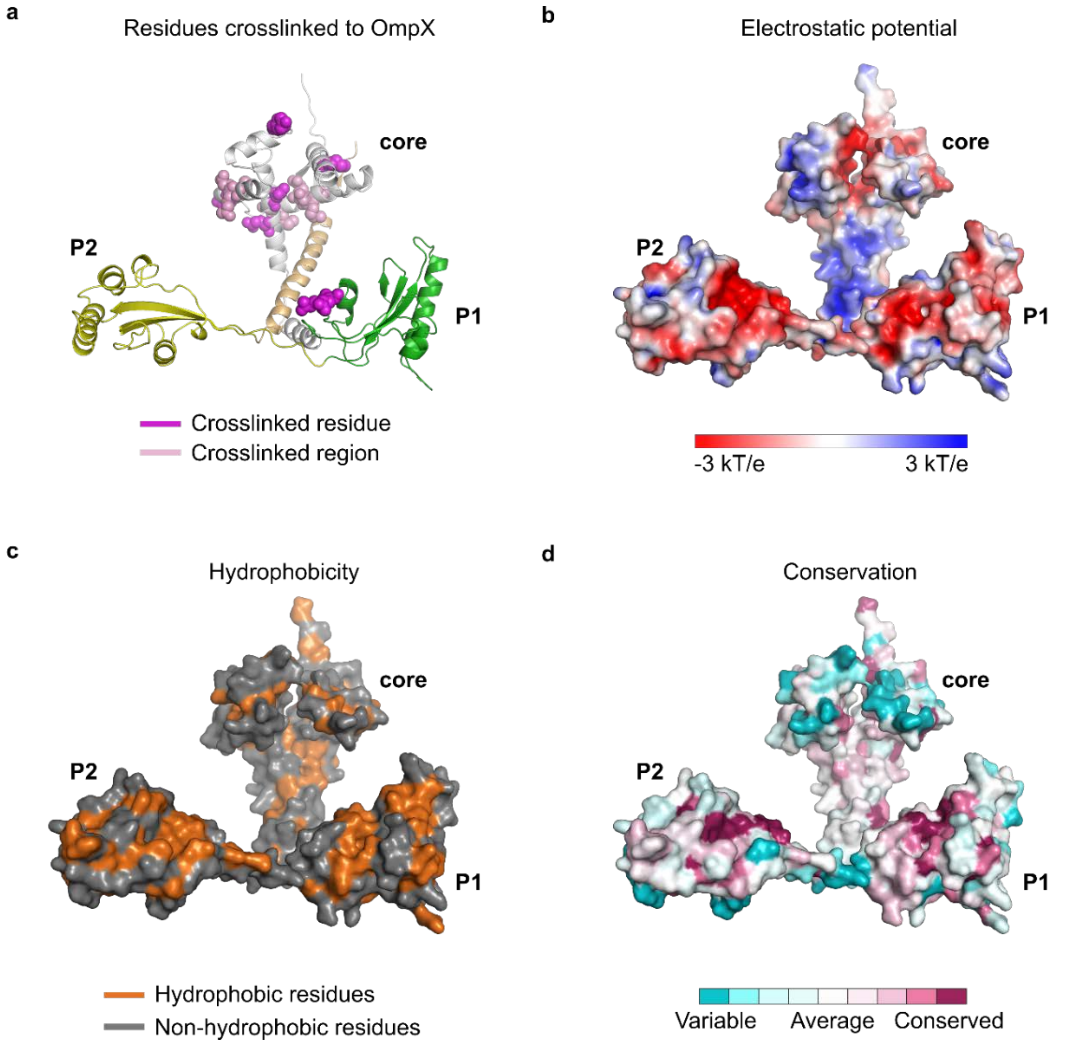


**Supplementary Fig. 13. Regions of SurA crosslinked to OmpX in tag-transfer crosslinking experiments show no obvious correlation with locations of a particular electrostatic surface potential, hydrophobicity or with areas of high conservation on SurA. (a)** Crosslinks identified in tag-transfer experiments between OmpX and SurA (**Fig. 5, Supplementary Fig. 9)** mapped to a single residue (dark purple) or short peptide (light purple). **(b)** Electrostatic surface representation of SurA (-3 kT/e to +3 kT/e) generated using the APBS plugin for PyMOL [8]. **(c)** Surface hydrophobicity of SurA. Hydrophobic residues (Gly, Ala, Val, Leu, Ile, Pro, Phe, Met, and Trp) are shown in orange, all other residues are highlighted in grey. **(d)** Residue conservation of SurA. Amino acid conservation is indicated as a colour gradient between cyan and red for variable and conserved residues, respectively. Conservation scores were generated using the ConSurf webserver (see Methods) [9]. Data are illustrated using the SurA$^{core\text{-}P1\text{-}open}$ model used as a starting structure for simulation (**Supplementary Fig.6a**).

**Supplementary Table 1: Inter-domain and intra-domain crosslinks identified for apo-SurA and their SLDs and SASDs in the crystal structure of *E. coli* SurA (PDB 1M5Y [2]).** DSBU has been shown to crosslink residues within ~27-30 Å ($C\alpha$-$C\alpha$ SLDs) [10]. To account for uncertainty due to the resolution in the crystal structure, crosslinks are defined as satisfied (blue) or violated (red) if they are separated by SLD of less than or greater than 28 Å, respectively. Crosslinks between residues with a $C\alpha$-$C\alpha$ SASDs of less than or greater than 35 Å are defined as satisfied (blue) or violated (red), respectively. An asterisk (*) indicates that the distance indicated is between a residue pair in which one partner is not present in the crystal structure and the distance is therefore an approximation based on filled missing loop residues using MODELLER [11] (see Methods).

| | Cross-links identified | Cα-Cα SLD (Å) | SASD (Å) |
|---|---|---|---|
| **Inter-domain crosslinks** | | | |
| Core-P1 | K251-K405 | 27 | 36 |
| | K252-K394 | ~26* | ~44 * |
| | K269-K394 | ~25* | ~46* |
| Core-P2 | K105-K278 | 28 | 32 |
| | K105-K293 | 51 | 55 |
| | K134-K278 | 41 | 53 |
| | K134-K293 | 67 | 80 |
| | K278-K394 | ~27* | ~32* |
| | K278-K405 | 29 | 37 |
| | K293-K394 | 54* | 71* |
| | K293-K405 | 60 | 79 |
| | K362-K405 | 50 | 58 |
| P1-P2 | K251-K278 | 21 | 27 |
| | K251-K293 | 63 | 73 |
| | K252-K278 | 21 | 26 |
| | K252-K293 | 63 | 76 |
| | K252-K362 | 47 | 53 |
| | K269-K278 | 26 | 36 |
| | K269-K293 | 66 | 84 |
| **Intra-domain cross-links** | | | |
| Core-Core | K86 K134 | 14 | 21 |
| | K90 K134 | 10 | 39 |
| | K134 K394 | ~26* | ~36* |
| | K134 K405 | 26 | 44 |
| | K394 K405 | ~16* | ~34* |
| P1-P1 | K252 K269 | 9 | 16 |
| P2-P2 | K278-K293 | 42 | 52 |
| | K278-K362 | 27 | 30 |
| | K278-K388 | ~16* | ~17* |
| | K293-K306 | 11 | 26 |
| | K306-K318 | 15 | 22 |
| | K315-K362 | 15 | 20 |
| | K362-K388 | ~34* | ~52* |

**Supplementary Table 2. Steady state anisotropy measurements.** Single cysteine variants of SurA were labelled with either Alexa Fluor 488 or Alexa Fluor 594 dyes and the anisotropy measured as described in the Methods. The measured steady-state anisotropies of all the samples are similar and low, indicating considerable dye mobility, allowing changes in $E_{FRET}$ to be ascribed to changes in inter-domain distances. Data are shown as mean ± SEM of three replicate measurements.

| Labelled SurA variant | Anisotropy |
|---|---|
| 488-Q85C | 0.14 ± 0.01 |
| 488-N193C | 0.16 ± 0.01 |
| 488-E301C | 0.16 ± 0.01 |
| 594-Q85C | 0.11 ± 0.04 |
| 594-N193C | 0.11 ± 0.08 |
| 594-E301C | 0.16 ± 0.06 |

**Supplementary Table 3. Inter-domain Lys-Lys crosslinks identified for apo-SurA and their minimum SLDs in the structures at the end of the three 1 μs simulations.** DSBU has been shown to crosslink residues within ~27 - 30 Å (Cα-Cα SLDs) [10]. To account for uncertainty due to the resolution the crystal structure, crosslinks are defined satisfied (blue) if the Cα-Cα SLDs of the residues involved are less than/equal to 28 Å, or violated (red) if they are greater than 28 Å. An asterisk (*) indicates that the measured distance is between a residue pair in which one partner is not present in the crystal structure and this distance is therefore an approximation based on filled missing loop residues using MODELLER [11]. Rep: repeat simulation. Note that SLDs were used here to assess compatibility with the crosslinks observed as calculating SASDs across the entire MD trajectories is not feasible computationally.

| Inter-domain crosslinks | Cross-links identified | 1M5Y distances (Å) | Distance (SLD) in the final structure of the simulation (Å) | | |
|---|---|---|---|---|---|
| | | | Rep 1 | Rep 2 | Rep 3 |
| Core-P1 | K251-K405 | 27 | 21 | 39 | 38 |
| | K252-K394 | ~26* | 25 | 33 | 34 |
| | K269-K394 | ~25* | 26 | 37 | 39 |
| Core-P2 | K105-K278 | 28 | 23 | 48 | 27 |
| | K105-K293 | 51 | 11 | 14 | 49 |
| | K134-K278 | 41 | 36 | 48 | 46 |
| | K134-K293 | 67 | 22 | 22 | 69 |
| | K278-K394 | ~27* | 18 | 18 | 28 |
| | K278-K405 | 29 | 23 | 29 | 39 |
| | K293-K394 | 54* | 27 | 33 | 45 |
| | K293-K405 | 60 | 28 | 29 | 59 |
| | K362-K405 | 49 | 36 | 37 | 41 |
| P1-P2 | K251-K278 | 21 | 13 | 14 | 22 |
| | K251-K293 | 63 | 36 | 43 | 52 |
| | K252-K278 | 21 | 16 | 16 | 19 |
| | K252-K293 | 63 | 39 | 47 | 49 |
| | K252-K362 | 47 | 38 | 39 | 32 |
| | K269-K278 | 26 | 21 | 20 | 25 |
| | K269-K293 | 66 | 40 | 49 | 55 |

**Supplementary Table 4: Inter-domain Lys-Lys crosslinks identified for apo-SurA and their minimum SLDs in MD simulations starting from a SurA$^{core\text{-}P1\text{-}open}$ model.** DSBU has been shown to crosslink residues within ~27 - 30 Å (Cα-Cα SLDs) [10]. To account for uncertainty due to the resolution in the crystal structure, crosslinks are defined satisfied (blue) or violated (red) if they are less than or greater than 28 Å, respectively. An asterisk (*) indicates that the measured distance is between a residue pair in which one partner is not present in the crystal structure and this distance is therefore an approximation based on filled missing loop residues using MODELLER [11]. Rep: repeat simulation.

| Inter-domain crosslinks | Cross-links identified | 1M5Y distances (Å) | Minimum distance (SLD) in the MD simulations (Å) | | |
|---|---|---|---|---|---|
| | | | Rep 1 | Rep 2 | Rep 3 |
| Core-P1 | K251-K405 | 27 | 18 | 15 | 19 |
| | K252-K394 | ~26* | 20 | 13 | 21 |
| | K269-K394 | ~25* | 21 | 18 | 29 |
| Core-P2 | K105-K278 | 28 | 21 | 31 | 12 |
| | K105-K293 | 51 | 9 | 8 | 17 |
| | K134-K278 | 41 | 30 | 45 | 32 |
| | K134-K293 | 67 | 18 | 16 | 24 |
| | K278-K394 | ~27* | 15 | 12 | 12 |
| | K278-K405 | 29 | 21 | 25 | 20 |
| | K293-K394 | 54* | 26 | 23 | 32 |
| | K293-K405 | 60 | 24 | 20 | 33 |
| | K362-K405 | 49 | 27 | 32 | 22 |
| P1-P2 | K251-K278 | 21 | 6 | 8 | 10 |
| | K251-K293 | 63 | 29 | 33 | 19 |
| | K252-K278 | 21 | 6 | 9 | 8 |
| | K252-K293 | 63 | 30 | 35 | 17 |
| | K252-K362 | 47 | 14 | 25 | 21 |
| | K269-K278 | 26 | 14 | 17 | 16 |
| | K269-K293 | 66 | 30 | 37 | 22 |

**Supplementary Table 5. Inter-domain Lys-Lys crosslinks identified for apo-SurA and their SLDs in the 10 lowest energy structures from simulated annealing MD.** Inter-domain crosslinks identified for apo-SurA and their SLDs in the crystal structure of *E. coli* SurA (PDB 1M5Y [2]) and in the 10 lowest energy structures from simulated annealing MD (**Supplementary Fig. 7**). Crosslinks are defined satisfied (blue) or violated (red) if they are less than or greater than 28 Å, respectively. Where the crosslink is not satisfied in the structure the Cα-Cα SLD is shown.

| Inter-domain crosslinks | Cross-links identified | 1M5Y distances (Å) | SLD between residues if crosslink is not satisfied | | | | | | | | | |
|---|---|---|---|---|---|---|---|---|---|---|---|---|
| | | | 1 | 2 | 3 | 4 | 5 | 6 | 7 | 8 | 9 | 10 |
| Core-P1 | K251-K405 | 27 | | | | | | | | | | |
| | K252-K394 | ~26* | | | | | | | | | | |
| | K269-K394 | ~25* | | | | | | | | | | |
| Core-P2 | K105-K278 | 28 | | | | | | | | | | |
| | K105-K293 | 51 | | | | | | | | | | |
| | K134-K278 | 41 | | | | 29 | | | | 31 | | |
| | K134-K293 | 67 | | | | | | | | | | |
| | K278-K394 | ~27* | | | | | | | | | | |
| | K278-K405 | 29 | | | | | | | | | | |
| | K293-K394 | 54* | | 30 | | | 31 | 33 | | 29 | | |
| | K293-K405 | 60 | | | | | 30 | 32 | | | | |
| | K362-K405 | 49 | | | 34 | 35 | 30 | | 35 | 30 | | 36 |
| P1-P2 | K251-K278 | 21 | | | | | | | | | | |
| | K251-K293 | 63 | | | | | | | | | | |
| | K252-K278 | 21 | | | | | | | | | | |
| | K252-K293 | 63 | | | | | | | | | | |
| | K252-K362 | 47 | | | | | | | | 30 | | |
| | K269-K278 | 26 | | | | | | | | | | |
| | K269-K293 | 66 | | | | | | | | | 32 | |

**Supplementary Table 6. Comparison of smFRET data and the 10 lowest energy structures from simulated annealing MD.** Table of the mean $E_{FRET}$ values for the three FRET pairs predicted from the 10 lowest energy structures from simulated annealing MD (**Supplementary Fig. 7**) and those from the smFRET experimental data (**Fig. 3**). Note that the smFRET distributions are broad and only the peak top values are shown.

| | Predicted $E_{FRET}$ | | | | | | | | | | |
|---|---|---|---|---|---|---|---|---|---|---|---|
| **Model** | **1** | **2** | **3** | **4** | **5** | **6** | **7** | **8** | **9** | **10** | **Experimental Data** |
| **Core-P1** | 0.23 | 0.12 | 0.12 | 0.1 | 0.1 | 0.09 | 0.1 | 0.5 | 0.85 | 0.1 | 0.6, 0.2 |
| **Core-P2** | 0.9 | 0.85 | 0.41 | 0.91 | 0.37 | 0.51 | 0.42 | 0.65 | 0.82 | 0.75 | 0.4 |
| **P1-P2** | 0.9 | 0.34 | 0.45 | 0.17 | 0.97 | 0.55 | 0.89 | 0.44 | 0.25 | 0.3 | 0.3 |

**Supplementary Table 7: Crosslinks detected between SurA and OmpX.** List of detected residues that were crosslinked with DSBU and the corresponding domains in SurA.

| Domain | SurA residue | OmpX residue |
|---|---|---|
| N | 87 | 50 |
| | 87 | 82 |
| | 91 | 82 |
| | 106 | 82 |
| | 135 | 50 |
| | 135 | 71 |
| | 135 | 82 |
| P1 | 253 | 71 |
| | 253 | 82 |
| P2 | 279 | 50 |
| | 279 | 82 |
| | 294 | 50 |
| | 294 | 71 |
| | 294 | 82 |
| | 307 | 50 |
| | 307 | 82 |
| | 316 | 82 |
| C | 389 | 50 |
| | 389 | 71 |
| | 389 | 82 |
| | 395 | 50 |
| | 395 | 71 |
| | 395 | 82 |
| | 406 | 71 |
| | 406 | 82 |
| | 425 | 82 |

**Supplementary Table 8: Modified SurA peptides identified from photocrosslinking experiments between SurA and MTS-diazirine-conjugated single Cys OmpX variants.** SurA residues found to be modified are shown in red. Where the spectral quality of the MS/MS spectra was insufficient to conclusively assign the modified residue, the residues to which the modification could be localised are shown in red.

| Sequence | Start Residue | End Residue | Modified Residue | OmpX Cys-variant |
|---|---|---|---|---|
| VAAVVNNGVVLESDVDGLMQSV | 28 | 49 | M46 | M41C, I102C, K122C, V167C |
| LIMDQIILQMGQK | 74 | 86 | I75 | M41C |
| IMDQIILQMGQK | 75 | 86 | I75 | M41C, I102C |
| IMDQIILQMGQK | 75 | 86 | M76 | M41C, I102C, V167C |
| LIMDQIILQMGQ | 74 | 85 | M83-G84 | M41C, I102C, V167C |
| IMDQIILQMGQ | 75 | 85 | I75-Q78 | K122C, V167C |
| SDEQLDQAIANIAK | 92 | 105 | S92 | I102C |
| SDEQLDQAIANIAK | 92 | 105 | L96-Q98 | V167C |
| LAYDGLNYNTYR | 118 | 129 | Y120 | M41C, M41C*, I102C, K122C* |
| EMIISEVR | 135 | 142 | E135 | M41C*, I102C*, V167C* |
| EMIISEVR | 135 | 142 | E140 | K122C*, V167C* |
| IQELPGIFAQALSTAK | 236 | 251 | E238 | M41C |
| QELPGIFAQALSTAK | 237 | 251 | Q237 | K122C |

* denotes identification in a sample enriched using thiopropyl Sepharose 6B resin

**Supplementary Movie Legends**

**Supplementary Movies 1-3: Three repeat MD simulations of full-length SurA using a SurA$^{core\text{-}P1\text{-}open}$ model as the starting structure.** 1 μs all-atom simulations of the mature sequence of SurA (residues 21-428) in explicit solvent were performed with GROMACS 5.0.2 [88] using the CHARMM36 force field [89]. The SurA$^{core\text{-}P1\text{-}open}$ model was built using the crystal structures of full-length SurA (PDB: 1M5Y [27]) and SurA-ΔP2 (PDB: 2PV3 [50]) in which the P1 domain is extended away from the core (see Methods). The N-terminal region of the core domain, P1, P2 and the C-terminal region of the core domain are shown in grey, green, yellow, and orange, respectively.
